# Predicting space use patterns of a territorial top predator: from individual movement decisions to Arctic fox space use

**DOI:** 10.1101/2025.09.29.679328

**Authors:** Frédéric Dulude-de Broin, Dominique Berteaux, Joël Bêty, Catherine Villeneuve, Alexis Grenier-Potvin, Andréanne Beardsell, Jeanne Clermont, Audrey Durand, Pierre Legagneux

## Abstract

1. Predicting animal space use could greatly improve our understanding and forecasting of ecological processes. Despite growing interest, the development of predictive space use models amenable to the integration of spatial processes into ecological frameworks have yet to reach their full potential.
2. Using high-resolution tracking data collected at 4-minute intervals from 26 Arctic foxes over five years, we developed a predictive space use model based on a step-selection approach. We assessed fine-scale habitat selection in relation to prey distribution, landscape features, and ecological constraints such as central place foraging and territoriality. We then used these results to build an agent-based model simulating fox space use and evaluated its ability to reproduce observed space use patterns.
3. Step-selection analyses confirmed that fox movements were driven by habitat type, goose nest density, distance to den, and avoidance of distance to the home range boundary. Agent-based simulations closely matched empirical tracking data and accurately forecasted fox space use, even for individuals excluded from model parameterization.
4. By developing a predictive model of predator space-use, our study provides a foundation for incorporating additional components of the predation sequence and contributes to more spatially informed approaches in predator-prey ecology.

## Introduction

Animal movement is fundamental to ecology. It shapes when, where and how frequently individual organisms interact, and ties landscape features to population dynamics and community structure (Beltran et al., 2024; Costa-Pereira et al., 2022). Incorporating spatial dynamics in population and community modelling could enhance our ability to understand and predict ecological processes (Beltran et al., 2024; Cherif et al., 2024; Jeltsch et al., 2013), but current approaches frequently assume well-mixed populations in which parameters are defined as functions of overall density (Fortin et al., 2015; Zurell et al., 2022). Well-mixed populations are rare in natural systems, as resources and consumers are typically structured in space and time by processes such as territoriality, grouping behaviour, or environmental heterogeneity. Yet, this assumption potentially results from the difficulty to account for the complex and highly stochastic nature of movement in modeling and analytical efforts (Nathan et al., 2008; Patterson et al., 2008), and the difficulty of collecting detailed high-resolution, tracking data from animals (Allan et al., 2018). Furthermore, linking individual movement decisions to population level pattern of space use is challenging because the movement process itself can generate emergent patterns (Potts, Bastille-Rousseau, et al., 2014).

Recent expansion of high-resolution spatial data (Nathan et al., 2022; Neumann et al., 2015; Williams et al., 2020; Wilmers et al., 2015) and analytical methods (Moorcroft et al., 2017; Potts & Börger, 2023; Smouse et al., 2010; Winter et al., 2024) now allows to model space use at scales that are relevant to study species interactions (Kays et al., 2015; Nathan et al., 2022). Such developments offer an unprecedented opportunity to integrate animal movement within population and community models.

In heterogeneous landscapes, animals should adopt movement behaviours that maximise food acquisition while navigating around physical barriers and minimizing energetic costs or risk from predators (Stephens & Krebs, 1986). Intraspecific interactions may further shape space use patterns by imposing constraints on habitat availability (Brown & Orians, 1970; Fretwell & Lucas, 1969; Grenier-Potvin et al., 2021). For instance, the monopolization of resources by some individuals may prevent others from accessing high-quality habitats, and reproduction can create spatial anchors (e.g. nest, den, spawning ground) impacting movement (Benhamou & Courbin, 2023). Drivers of space use are often assessed through resource selection functions (RSF), which compare habitat features at used and available locations using logistic equations (Manly, 2002). RSF enabled valuable progress in the field of movement ecology, but do not explicitly consider animal movement capacity and assume resources are equally accessible to individuals, a premise rarely verified in natural settings (Buskirk & Millspaugh, 2006). Consequently, RSF methods are typically limited to relatively coarse scales where data resolution is low enough to reduce autocorrelation issues (Fieberg et al., 2020). In addition, habitat complexity or physical impediments such as roads, rivers, and patches of inhospitable habitat can obstruct animal movement (Beyer et al., 2016; Smith, Donadio, Pauli, Sheriff, Bidder, et al., 2019), but are generally omitted in RSF overlooking their impact on space use (Potts, Bastille-Rousseau, et al., 2014). On the other hand, step-selection functions (SSF) are another set of habitat selection analyses that solve some of the above-mentioned RSF limitations. Notably, SSF use a conditional logistic equation to define habitat availability for each segment between consecutive relocations, allowing joint estimation of movement capacity and habitat selection (Avgar et al., 2016; Fortin et al., 2005). Predictions of space use can then be made using differential equations or by iteratively simulating the relocation of individual agents based on empirically derived selection coefficients (Potts, Bastille-Rousseau, et al., 2014; Potts & Börger, 2023; Signer et al., 2024). While SSF hold great promises for forecasting animal movement and interest in their application is growing, they have not yet been widely applied to build predictive models in natural settings (Gomez et al., 2025; Potts & Börger, 2023).

The study of predator-prey interactions could greatly benefit from predictive space use models using movement data (Cherif et al., 2024). Spatial dynamics can play a key role in predator-prey interactions (Schmitz et al., 2017; Sih, 2005), and the behavioural response of individuals to landscape features can strongly alter their outcomes (Labadie et al., 2023; Oliver et al., 2009). At broad scale, behavioural trade-offs influence home range size and population densities (López-Sepulcre & Kokko, 2005; Loveridge et al., 2009; Sells et al., 2022), which in turn may affect prey survival (Dulude-de Broin et al., 2023) and species coexistence (Beardsell et al., 2023). At finer scale, differences in time budget between predator and prey (Gehr et al., 2018; Smith, Donadio, Pauli, Sheriff, & Middleton, 2019), variations in foraging efforts (Beardsell et al., 2022; Lichtenstein et al., 2019), and specific habitat preferences (Heithaus et al., 2009) may disconnect local predator abundance and realised encounters with prey. The incorporation of such spatial dynamics in predator-prey models is a natural next step in our understanding of community dynamics (Suraci et al., 2022). However, despite growing interest in forecasting animal space use (Aiello et al., 2023; Potts & Börger, 2023; Signer et al., 2024; Vanlandeghem et al., 2021), most movement studies still focus on identifying the drivers of animal movement rather than building predictive models (Beltran et al., 2024; Potts & Börger, 2023) .

Here, we used high-resolution tracking data on a terrestrial top predator to develop a predictive model of space use based on SSF approach. Building on the work from Grenier-Potvin et al. (2021) that used RSF to study Arctic foxes habitat preferences, we applied SSF to assess and predict fine-scale habitat selection in relation to prey distribution and landscape features during the bird incubation period in the Arctic tundra. Our model also incorporated the key ecological constraints of our system, including central place foraging and territoriality. We evaluate the model’s predictive performance by comparing simulated trajectories with observed movement data. This study contributes to the development of more spatially informed approaches in predator–prey ecology.

## Materials and Methods

### Study system

The study was conducted from 2018 to 2023 in the southwest plain of Bylot Island (72.93° N, 79.76° W), in Sirmilik National Park, Nunavut, Canada. The landscape is primarily composed of low-elevation mesic tundra and high-center polygonal wetlands, interspersed with xeric tundra, wet meadows, gravel beds, as well as thaw lakes, rivers, and ponds. It includes a large snow goose (*Anser caerulescens*) colony, occupying ca. 70 km^2^ where around 5,000 to 20,000 pairs nest each year in a patchy distribution driven by snowmelt patterns during nest initiation (Reed et al., 2002).

Foxes mainly feed on brown and collared lemmings, as well as greater snow goose eggs when available. Lemmings follow marked cycles of abundance with a period of 3-4 years (Fauteux et al., 2015; Gruyer et al., 2008). Brown lemming, the most abundant species, is predominantly found in mesic tundra and heterogenous wetlands, while collared lemmings are most often distributed in mesic and xeric habitats (Szor et al., 2008). Goose eggs provide a major food source for foxes in the vicinity of the colony. Geese typically nest in high-center polygonal wetland and mesic tundra habitats. In addition, other bird species such as Cackling geese (*Branta hutchinsii*), Glaucous gull (*Larus hyperboreus*), Red-throated loon (*Gavia stellata*) and various shorebird and passerine species nest at low density across the island (Gauthier et al., 2011). These species at low densities were not included in analyses as they represent only a small portion of fox diet (Giroux et al., 2012) and should have minimal influence on fox movement.

### Fox movement data

Between May and August from 2018 to 2023, we captured 26 adult foxes within the snow goose colony using padded leghold traps (model Softcatch #1, Oneida Victor Inc. Ltd., USA). Foxes were marked with color ear-tags for visual identification and equipped with solar-powered GPS collars (95 g, ca. 3-4% of body mass; Radio Tag-14, Milsar, Romania). Some individuals were tracked across multiple years, yielding a total of 39 fox-years of movement data. GPS locations were collected every 4 minutes. For this study, we used data collected during the goose nesting period defined annually by the average laying and hatching dates in the goose colony. All individuals included in the analyses had access to the goose colony during this period.

### Behavioural state (HMM)

We used a Hidden Markov Model implemented through the moveHMM R package (Michelot et al., 2018) to classify movement data in two behavioral states: active (characterized by long step lengths and small turning angles) and inactive (characterized by short step lengths and large turning angles). Time-of-day was included as covariate to account for fox circadian rhythm. The Hidden Markov Model performed well and allowed for the identification of the two hypothetical behavioral states (Grenier-Potvin et al., 2021).

### Movement covariates Habitat class

We used the habitat map created by Grenier-Potvin et al. (2021), which was based on a 0.5-m resolution WorldView-2 satellite image from July 2, 2018, and generated using random forests with a hybrid object-based approach (Chen et al., 2017). This supervised method, validated through ground-truthing, provided high classification accuracies for distinguishing water/ice/snow from land (98.6%), classifying land cover types including mesic, wet meadow, xeric, gravel and sand (89.4%), and identifying polygonal wetlands (93.3%) (Grenier-Potvin et al., 2021).

### Goose nest density

In 2018 and 2019, a proxy for goose nest density was mapped across the study area through field surveys on foot to assess the relative spatial availability of this abundant prey item. Nesting geese were counted within homogeneous patches, which were delineated and georeferenced using prominent landmarks such as lakes, rivers, and rocks. The relative goose nest density derived from this method showed strong agreement with systematic nest counts conducted in random plots (Grenier-Potvin et al., 2021). For 2018 and 2019, we thus had a layer of goose nest density to inform prey availability for foxes. Similar goose count surveys were not conducted in subsequent years. However, the location and extent of the goose colony are generally consistent across years (Duchesne et al., 2021), and a pixel-wise comparison of 2018 and 2019 data revealed strong among-year correlation in goose nest density (Spearman’s r = 0.75 [0.74–0.76], n = 18086). For years after 2019, we thus mapped spatial patterns of relative goose nest density by averaging values from the 2018 and 2019 maps. We also confirmed that goose nest density coefficients were similar whether the SSF model was fitted to all years or to 2018–2019 only.

### Fox home range estimation

Foxes within the study area have stable and largely exclusive home ranges (Clermont et al., 2025). To include this behaviour in our analyses, we determined the annual home range of foxes (95% isopleth) through autocorrelated kernel density estimation (Fleming et al., 2015), using the ctmm R package (Calabrese et al., 2016). Following Fleming et al. (2015), we created scatter plots of the relocation data and calculated empirical variograms to identify and exclude any extraterritorial excursions and ensure that foxes were range resident (i.e. remained within a spatially confined area). We then selected an appropriate home range model for each individual by comparing different movement processes (independent identically distributed, Ornstein–Uhlenbeck, integrated Ornstein–Uhlenbeck, and Ornstein–Uhlenbeck Foraging) using the Akaike Information Criterion (AICc), as described by Calabrese et al. (2016). The area of the 95% home range contour was estimated for each individual-year based on the selected model.

### Reproductive status and main den location

Reproductive status and main den location of marked individuals were determined through camera trap monitoring of known dens within the study area (Cameron et al., 2011). For breeding individuals, the natal den was considered the main den. For non-breeding individuals, we used the den with the highest activity.

### Step-selection function

We assessed fox movement preferences by fitting an integrated step-selection function (SSF; Avgar et al., 2016; Fortin et al., 2005) on data collected between 2018 and 2023 (n=31 fox-year), leaving out data from 25% of the individuals using stratified random sampling (8 foxes: 2 from 2018, 3 from 2019, 2 from 2022, and 1 from 2023) to further assess the predictive performances of our simulations. SSFs are discrete choice models that compare environmental features and movement characteristics of observed steps (the linear segment between consecutive relocations) to alternative random steps originating from the same location. We performed the SSF on continuous sequences (bursts) of relocations every 4 minutes with a tolerance of ±30 seconds, keeping only active locations as identified by the Hidden Markov Model. For each observed step, we generated 50 random steps by sampling step lengths and turning angles from empirical gamma and Von Mises distributions, fitted to the original movement data. Covariates were then extracted by taking the average value of random points sampled every two meters along the entire linear segment defined by each step, rather than just at the endpoint. This method ensured that the simulated movement trajectories would reflect habitat preferences of foxes, penalizing steps that intersect with avoided habitats. Covariates included in the model were the proportion of each habitat type, the average value of goose density, log-transformed distance to home range edges, and log-transformed distance to den in interaction with reproductive status. These variables were identified as important drivers of arctic fox movements in Grenier-Potvin et al. (2021). Additionally, the model included step length, log-transformed step length, and cosine of turning angles as movement parameters. We fitted the SSFs using generalized linear mixed models with a Poisson distribution and a stratum-specific intercept implemented with glmmTMB package in R (Brooks, Mollie et al., 2017). This method allows the inclusion of random effects and is likelihood-equivalent to SSF models fitted using conditional logistic regression (Muff et al., 2020). To account for the grouping structure of the data and allow selection to vary across individuals and years, we included random slopes for each variable by individual-year (i.e. a unique identifier for each individual each year). The generation of predicted values with confidence intervals from the SSF model fitted with glmmTMB was challenging due to very large model size and heavy computation needed to calculate the variance-covariance matrix with random effects. For visualization purposes only, we therefore used predictions from the same model without random effects. This had no impact on results reported in text and simulations, which are all based on coefficients from the model including random effects.

### Movement simulations

We originally developed a movement simulator independently, using an approach that was later found to be similar to the simulate function in the newly released amt R package (Signer et al., 2024). After its release, we refined our simulator by integrating relevant features from their implementation, which improved computational efficiency of our simulator. Relocations were generated through an iterative process starting with the sampling of 100 random steps from the empirical distributions of step-lengths and turning angles, thus avoiding unrealistic movement steps. At each step, covariate values were extracted, and the relative selection probability (weight) was calculated based on the SSF coefficients. A step was then selected probabilistically, with selection probability proportional to its weight. Finally, the simulated fox position was updated to the endpoint of the chosen step. This process was repeated to generate the complete set of relocations. Relocations were only generated for fox active state, and the total number of relocations thus reflects the proportion of time spent active. To ensure that simulations remained within the home range boundaries, steps falling outside the home range were assigned a weight of 0. We simulated fox movements using the home range, den locations, goose nest density, and habitat map of all GPS-collared individuals. Simulations were conducted over 23 days, reflecting the snow goose incubation period. Daily fox activity was set to 50% based on fox activity data (Beardsell et al., 2022; Grenier-Potvin et al., 2021). We considered two scenarios: (1) a *SSF informed model*, in which movement decisions were guided by the selection coefficients estimated from the SSF model, and (2) a *Selection-free model,* in which no habitat selection was applied. In both cases, fox movements were constrained within their home ranges, and potential steps were drawn from the empirical distributions of step lengths and turning angles. The only difference between scenarios was therefore the use (or absence) of selection coefficients. We generated 10 simulations per scenario for each observed fox.

### Comparison between observed and simulated foxes

We assessed how well our model could predict fox movement patterns by comparing observed fox trajectories to simulated trajectories from the SSF-informed and Selection-free models. Simulations were first evaluated using all GPS-collared foxes including those that were used to fit the SSF model to maximize sample size and assess how well the model reproduces overall movement patterns. We then tested the model by comparing simulated and observed trajectories only for individuals that were withheld when fitting the SSF model to evaluate predictive performance beyond the original dataset.

Specifically, we compared tracks using the ratio of used to available locations for each covariate. To measure use, we randomly sampled locations every 10-meters along the trajectories of observed and simulated foxes and extracted i) habitat class, ii) goose density, iii) distance to den and iv) distance to home range border. To assess availability, we sampled 500 000 random points within each fox’s home range and extracted the same four metrics. Finally, for each covariate, we calculated the use to availability ratio. We then compared the relative use of habitat features using linear mixed models. A separate model was fitted for each covariate with relative use as the response variable, data type (three-level categorical: Observed, Selection-free, SSF-informed) as a fixed effect, and fox map ID as a random effect.

To compare overall space use patterns, beyond habitat covariates, we also derived and compared utilization distributions (UD) from predicted and observed fox tracks. UDs represent the probability of an animal using different parts of its range and can be visualized as heatmaps of relocation density. The resulting spatial patterns of utilization (i.e. heatmaps) from observed and simulated foxes were compared using Earth Mover’s Distance (EMD; Potts, Auger-Méthé, et al., 2014) implemented in the *transport* R package (Schuhmacher et al., 2024). EMD quantifies the dissimilarity between two distributions by calculating the minimum effort required to transform one into the other, with smaller values indicating greater similarity. Given the short duration of tracking and simulations, it is unlikely that either the observed or simulated trajectories reached stable utilization distributions (UDs), making the resulting EMD values sensitive to stochastic variation. While we expect substantial variability in EMD values, comparing the average distance of UDs derived from observed tracks to UDs derived from simulated tracks (SSF-informed and Selection-free) should offer an additional indicator of how well the model captures observed movement patterns.

## Results

Between 2018 and 2023, we tracked 39 Arctic foxes at 4-minute intervals during the goose incubation period, monitoring individuals for an average of 19 days (range: 5–22 days). Average home-range size was 12 km² (range: 3–30 km^2^), and goose nests were available within every home range. Once calibrated with the step-selection analysis, the movement model successfully generated fox movement trajectories based on den locations, habitat map, goose nest density and home range contours (Figure 1).

**Figure 1.**
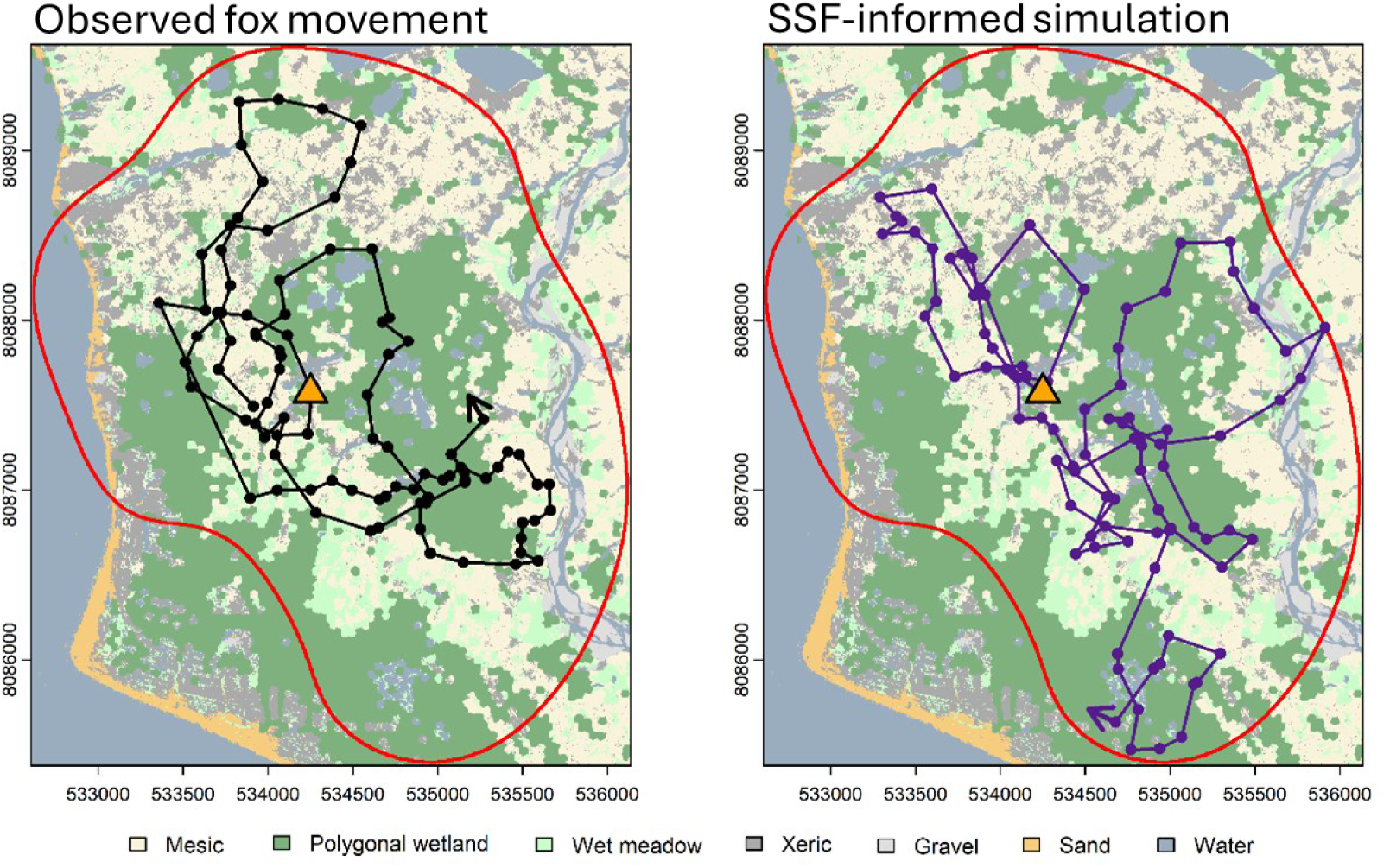
Example of 6-hour long simulated (left) or observed (right) fox trajectories on Bylot Island, Nunavut, Canada. The purple and black lines illustrate simulated and observed movement paths respectively with dots showing predicted and observed relocations at 4 minutes intervals. An arrow was added at the end of the trajectory. Trajectories are displayed over the habitat map, with the main den (orange triangle) and home range contour (red line).

Step-selection analyses confirmed that fox movements were driven by habitat type, goose density, distance to den, and distance to home range boundary (Fig. 2 and 3). Cross validation results indicated excellent model fit (spearman r [95%CI] = 0.94 [0.93,0.95]). Foxes strongly avoided gravel beds (Estimate[95%CI] = -0.13 [-0.15, - 0.10]) and water bodies (-0.18 [-0.22, -0.15]) compared to mesic habitat (the reference category; Fig. 2). They also slightly avoided wet meadows (-0.04[-0.07, -0.02]) and polygonal wetlands (-0.05[-0.08, -0.01]) and marginally preferred xeric habitat (0.06 [0.03, 0.09]), although effect sizes were much smaller for these three habitat categories (Fig. 2). Selection of sand habitat was neutral (0.01[-0.04, 0.05]). Step selection probability increased with snow goose nest density (0.17 [0.11, 0.23]; Fig. 3, top). Foxes also preferred steps that brought them closer to their den (-0.21 [-0.38, -0.04]) and the effect was stronger for reproductive individuals than for non-reproductive ones as indicated by the interaction term (-0.26 [-0.48, -0.03]); Fig. 3, middle). Foxes preferred steps that led them away from the home range boundary (0.81 [0.72, 0.90]), with negative selection observed for distances less than 400 meters from the edge (Fig. 3, bottom). Random slopes in the SSF model revealed substantial inter-individual variability for step length (step-length: SD = 0.50, log step-length: SD = 0.25), followed by proximity to den (log den: SD = 0.27), territory borders (log border: SD = 0.215), and goose nest density (SD = 0.15). Inter-individual variations in habitat selection coefficients were comparatively small (SD ≤ 0.103).

**Figure 2.**
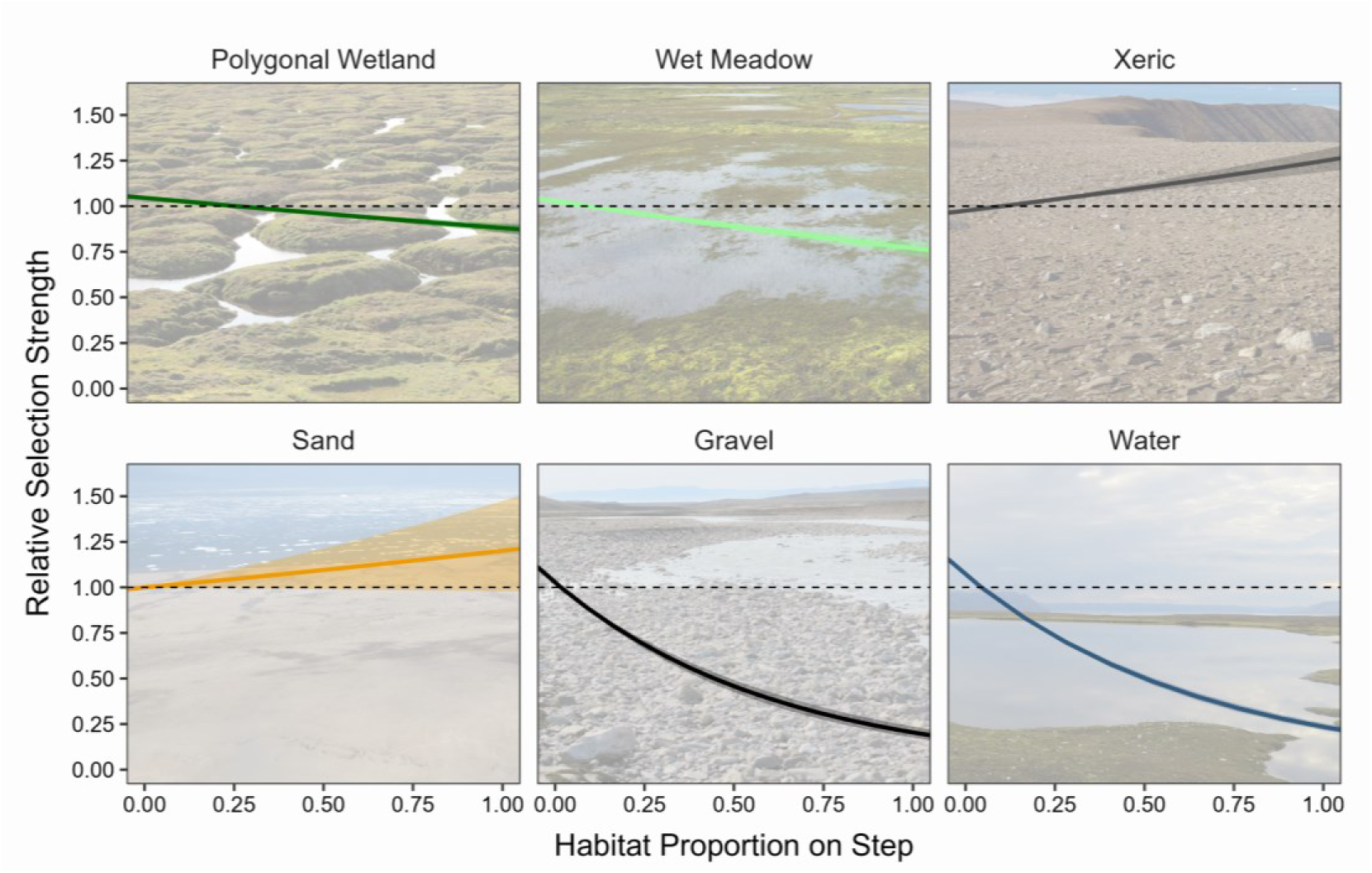
Effect of habitat proportion on step selection probability relative to mesic habitat based on the step-selection function fitted on 31 individual-year foxes equipped with GPS collars on Bylot Island, Canada, between 2018–2023. Each panel represents a different habitat type, with slopes indicating the relative selection strength and shaded areas showing 95% confidence intervals. Values above 1 indicate a higher selection probability compared to mesic habitat, whereas values below 1 indicate lower selection probability, with all other covariates held constant.

**Figure 3.**
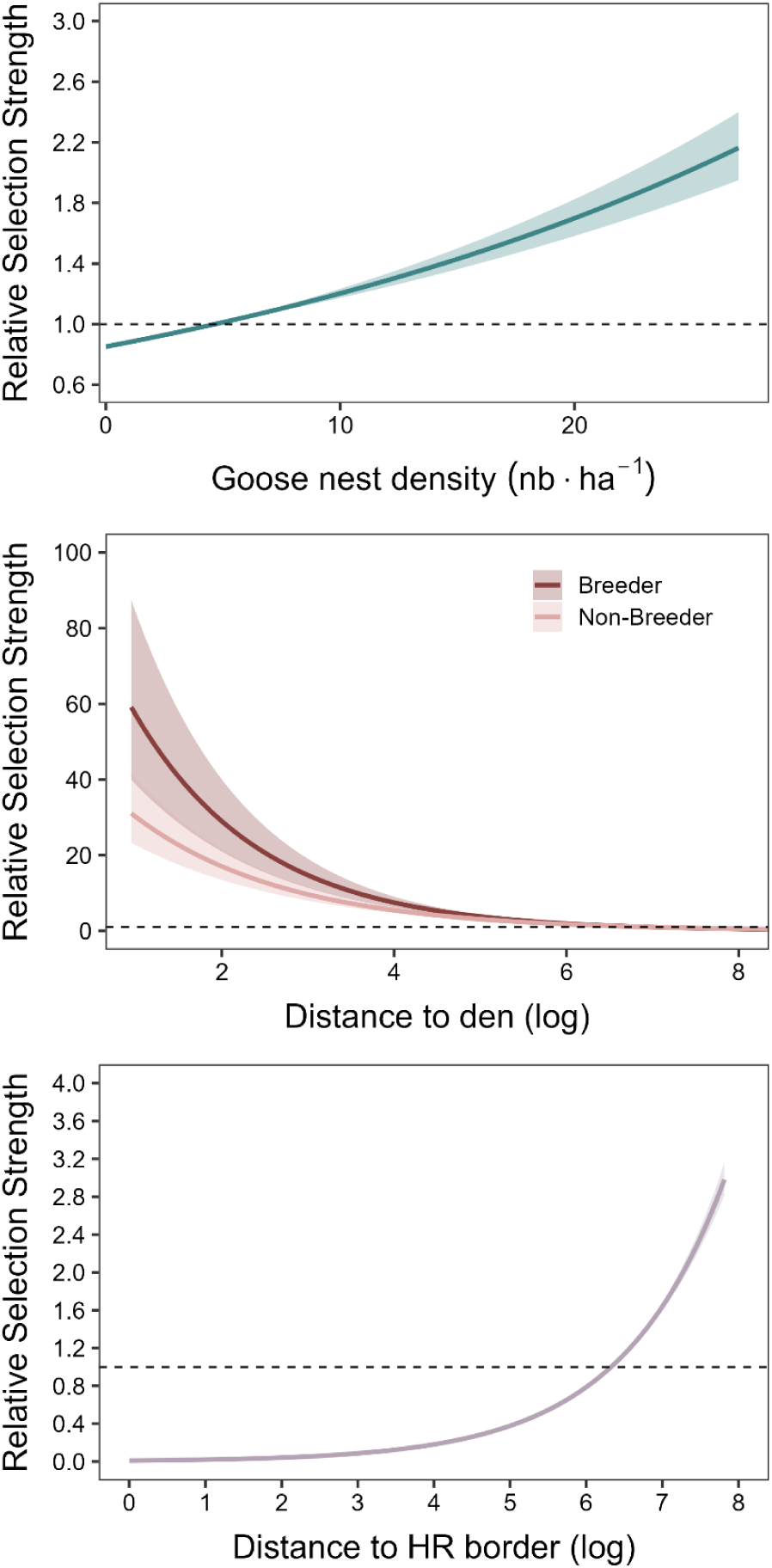
Effect of snow goose nest density (top), distance to den (middle), and distance to home range (HR) border (bottom) on step selection probability based on the step-selection function fitted on 31 individual-year foxes equipped with GPS collars on Bylot Island, Canada, between 2018–2023. Slopes represent relative selection strength compared to the mean and are shown with their 95% confidence intervals (shaded areas). Values above 1 indicate a higher selection probability than average, while values below 1 indicate a lower selection probability, with other covariates held constant.

The SSF-informed movement simulations were consistent with habitat classes visited by GPS-collared foxes (Figure 4). Across all habitat classes, relative use was closer between observed tracks and SSF-informed simulations than between observed tracks and selection-free simulations (see Appendix S1 for full model coefficients). The improvement was especially notable for sand, gravel, and water habitats (Figure 4; Appendix S1). For mesic, polygonal wetland, wet meadow, and xeric habitats, which had much weaker selection coefficients, the SSF-informed model still performed slightly better than the selection-free model but differences between models were minimal (Figure 4; Appendix S1). Interestingly, sand avoidance emerged in the simulations despite the neutral selection coefficient in the SSF (Figure 2 and 4), reflecting patterns generated by the movement process in unevenly distributed habitat patches.

**Figure 4.**
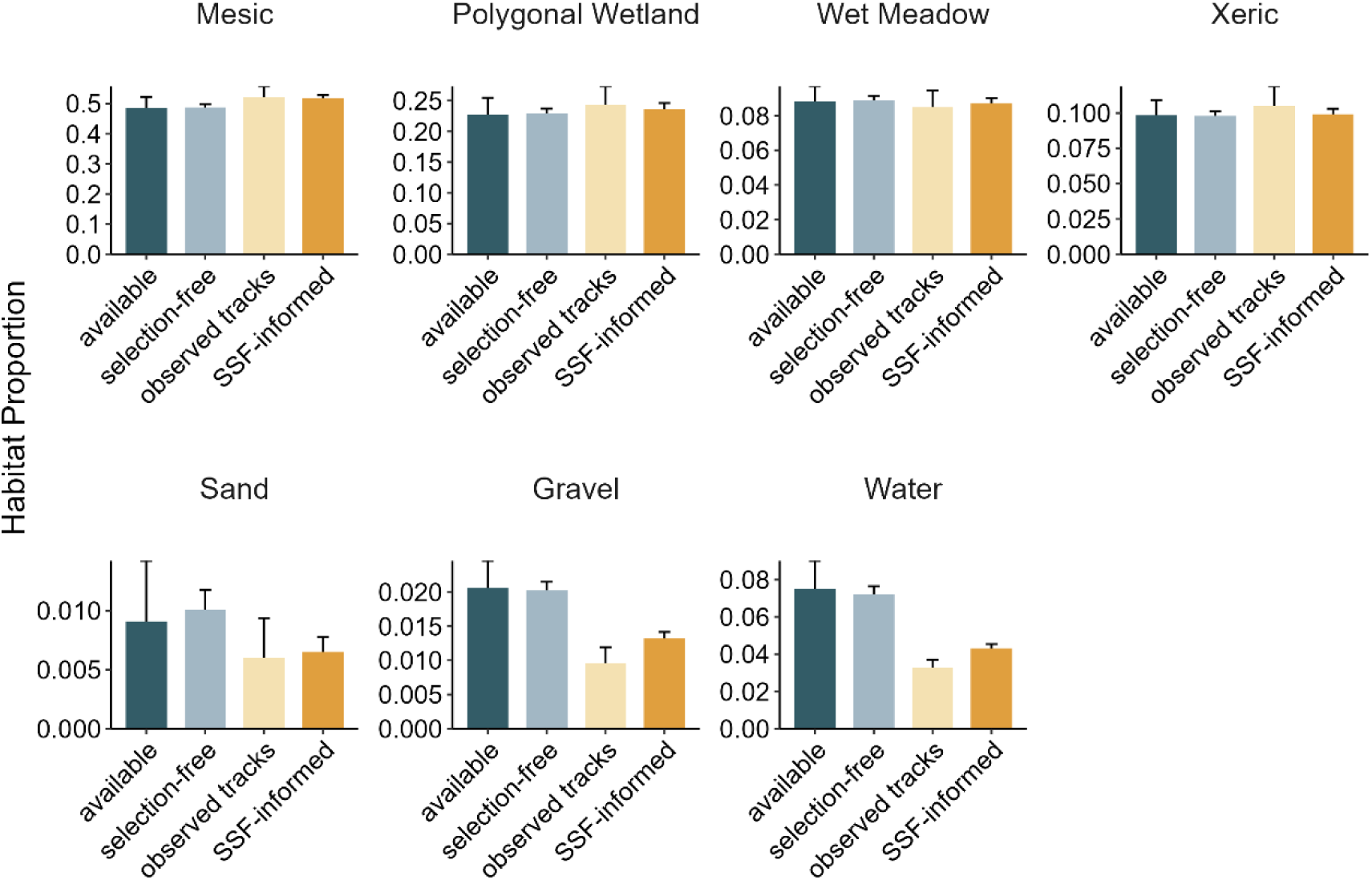
Comparison of average habitat class proportions across four categories: available (habitat available within fox home ranges), random (habitat visited by simulated tracks from a null movement model assuming no habitat selection), observed (habitat visited by GPS-collared foxes), and simulated (habitat visited by simulated tracks from the final movement model incorporating habitat selection). This comparison evaluates how closely habitat use by simulated fox tracks from the final model matches observed patterns on Bylot Island, Canada, between 2018 and 2023.

Patterns of habitat use in relation to goose nest density, distance to the den, and distance to the home range boundary were also consistent between observed and SSF-informed simulated trajectories (Figure 5). The SSF-informed model outperformed the selection-free model across all features and did not statistically differ from observed tracks for distance to the den and home range boundary (Figure 5; Appendix S1). However, although it also outperformed the selection-free model for goose nest density, the SSF-informed model underestimated the influence of this variable, particularly at high nest densities (Figure 5; Appendix S1). Step-length and turning angles drawn from the empirical distribution were similar across models and consistent with observed tracks (Appendix S1).

**Figure 5.**
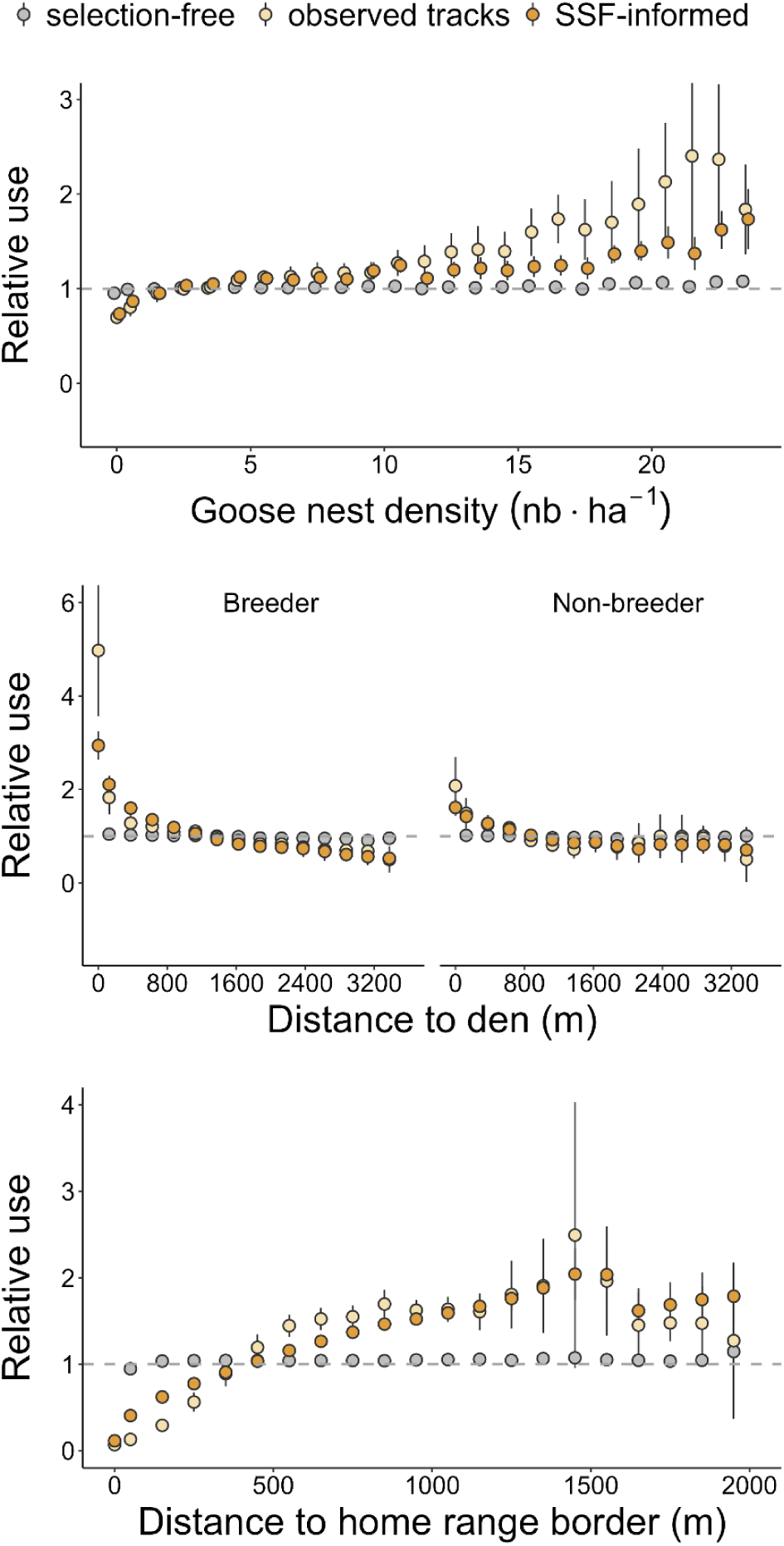
Comparison of the relative use of habitat patches with different goose densities, distance to den and distance to home range border for observed tracks (yellow), simulated tracks generated from the SSF-informed movement model (orange) and simulated tracks generated from a selection-free movement model (gray). Dots represent the relative use (i.e. the ratio of used to available habitat patches) within equally sized bins, averaged across all individuals or simulations, with error bars indicating 95% confidence intervals.

The comparison of overall utilisation distributions aligned with the analysis of specific habitat features. UDs from tracks generated using the SSF-informed model had on average lower dissimilarity to observed spatial utilization patterns (average Earth Mover’s Distance with observed tracks [95% CI]: 240.8 [226.1, 255.5]) than UDs derived from the selection-free model (average EMD [95%CI]: 302.7 [288.0, 317.4]).

Results on the restricted test sample (n=8 foxes withheld when fitting the initial SSF model) mirrored those obtained using all individuals, although the small sample size considerably increased estimates variability (see Appendix S1 for full results). Overall, the SSF-informed model clearly outperformed the selection-free model and showed high consistency with observed fox tracks. While there is still room for improvement, such as the model’s underestimation of use in areas of high goose nest density, our results demonstrate that it effectively captured fox movement patterns.

## Discussion

We developed a predictive model of space use by a top Arctic tundra predator using high-resolution tracking data, environmental covariates and prey distributions. The model was well adjusted to empirical tracking data and able to forecast fox movements, even for individuals that were not used to parametrize the model.

### Fox habitat preferences

The step-selection function confirmed findings on fox habitat preferences from Grenier-Potvin et al. (2021) based on resource selections functions. It highlighted strong avoidance of gravel and water, while other habitat class had smaller effects. The availability of goose nest density, a major prey for foxes, also had substantial influence on fox movements. Proximity to the den and distance to home range border were among the strongest drivers of movements, reflecting the importance of central place foraging and territoriality for this predator.

The inclusion of random effects into the step-selection model revealed substantial inter-individual and inter-annual variation in selection coefficients, particularly for step length. This is important as step-length is directly linked to daily distance travelled, a key parameter driving prey acquisition rates by foxes (Beardsell et al., 2022; Toscano & Griffen, 2014). Similarly, there was moderate variations in the level of attraction to the den, avoidance of border and selection of goose nest density among individual-years. We accounted for this variability when estimating SSF coefficients and based our simulator on population-level predictions, resulting in a generalizable model of fox movement. However, among-individual differences in behaviour of both predators and prey could influence the outcome of predator-prey interactions (LaBarge et al., 2024; Toscano et al., 2016). Considering these individual differences in predictive models could help refine our understanding of food web dynamics and should be the focus of future studies. For instance, it would be interesting to investigate how consistent individual variability in behaviour influence predator-prey dynamics (Hertel et al., 2020; LaBarge et al., 2024). Our fox space-use model could be used to study such dynamics by simulating varying levels of inter-individual differences in movement parameters and assessing their effects on prey encounter rates.

### Predicting space-use patterns

The SSF-informed model effectively captured the movement patterns of observed foxes and consistently outperformed the selection-free model. However, not all covariates contributed equally to fox space use. For instance, the effects of the most abundant habitat classes were weak. Indeed, although the SSF detected subtle differences in fox habitat preference for polygonal wetland, wet meadow and xeric habitat, the influence of these habitat class on overall fox space use was not clearly apparent in both the empirical and simulated tracks. This is likely due to the stronger effect of other variables such as snow goose nest density, distance to the den and distance to home range border which overshadowed habitat type. However, the avoidance of gravel and water were obvious in both the empirical and simulated tracks.

Our simulations illustrated that habitat structure itself, not just habitat type, can shape space use patterns. For example, sand avoidance emerged in the SSF-informed simulations despite the SSF selection coefficient for sand being neutral (Figures 2 and 4). This is likely due to the avoidance of home range boundaries where most sand patches were located (e.g. Figure 1). Similar effects could be expected when large patches of avoided habitat (e.g. water or gravel) fragment the landscape, or if selected features such as goose nest density or proximity to the den covary with other habitat features. These patterns underscore the value of more mechanistic approaches for generating space-use patterns, such as iterative individual-based simulations informed by SSF models. Indeed, emergent behaviours induced by habitat structure could be missed when generating tracks from resource selection functions or from predictions based solely on selection coefficients. This is especially limiting in cases where there is an interest in predicting space use for new habitat configurations that were not seen when assessing habitat preferences.

The effect of goose nest density on fox space use was clear, but appeared underestimated in the simulations compared to observed foxes. This could be due to memory driven processes (Fagan et al., 2013), as observed foxes are likely able to remember and travel to areas with high prey concentration, whereas simulated foxes did not have this ability. Indeed, our SSF-informed model could only predict local short-term movement decisions and was not driven by long-term goals. Incorporating hierarchical movement decisions at different scales could help better capture memory-driven processes and spatial behaviour in relation to clustered resources (Fagan et al., 2013). For example, combining a model that predicts fox movement direction at a broader scale (e.g. 30 minutes steps) with one that captures short-term movement decisions (e.g. 4 minutes steps) might better reflect space use in relation to goose nest density.

In our simulations, we imposed few strict rules and the predictive model was entirely calibrated from empirical SSF coefficients. The only notable exception is that we constrained simulations within defined home ranges. Home range size is shaped by complex behavioural processes likely involving trade-offs between the availability of food resources, competition from conspecifics, and individual competitive abilities (Sells & Mitchell, 2020). Our study was focused on modelling fox movement within existing home ranges, and we lacked the information needed for robust home range boundaries to emerge from the simulations. However, developing mechanistic models of home range size (Potts & Lewis, 2014) would be a natural next step to capture fox space use at broader scales.

### Implications for predator-prey interactions

To properly evaluate the outcome of habitat preferences on predator space use, relative habitat effects must be interpreted in relation to other covariates. In our system, the dominant habitat classes (mesic, polygonal wetland, xeric and wet meadow) had only a weak influence on predator space use and their effect was negligible compared to those of goose nest density, proximity to den and home range boundaries. Predator-prey encounter rates are thus likely more influenced by the availability of alternative prey species and interspecific interaction than by habitat class. The only clear exceptions were gravel beds, which foxes strongly avoided, and sand habitats, which occurred primarily along avoided home range edges.

Positive habitat-mediated effects of polygonal wetlands for snow geese (Lecomte et al., 2008) and gravel beds for common ringed plover (*Charadrius hiaticula*; Léandri-Breton & Bêty, 2020) have been reported at our study site. Our results confirm that fox strong avoidance of gravel translates into lower time spent in these habitats, which should reduce encounter rates to the benefit of ringed plovers. Similarly, empirical studies have also reported higher predation risk in areas with high goose nesting density (Mckinnon et al., 2013), which is again consistent with the higher predicted utilisation of these patches by foxes. However, observed and simulated tracks suggest that fox weak avoidance of polygonal wetlands is unlikely to account for the positive effect on snow goose nesting success, as there was no clear reduction in time spent within this habitat. Instead, other behavioural processes such as differences in attack rates or hunting success may better account for the observed pattern. Our results show that changes in encounter rates due to fox habitat preferences can explain observed predation patterns (e.g. higher nesting success in gravel beds, lower in patches of high goose density) but also suggest that other mechanisms are at play (e.g. higher nesting success in polygonal wetland despite high utilisation by foxes). To further explore these mechanisms, additional stages of the predation sequence beyond spatial avoidance (detection, attack, success probabilities) could be incorporated (Beardsell et al., 2021; Wootton et al., 2023).

The model we present allowed to predict fox space use patterns but did not explicitly incorporate temporal dynamics apart from the proportion of time spent active. Spatio-temporal variations of risk can be important in some systems as prey may benefit from predator downtime to access areas that are profitable but risky when predators are active (Dröge et al., 2017; Palmer et al., 2022). In the Arctic tundra, the primary prey of foxes are spatially anchored to their nests or burrows. Their spatial domain is small compared to that of foxes which may limit their ability to dynamically mitigate risk through spatial avoidance (Schmitz et al., 2017). However, most bird species take incubation recesses and lemmings often forage outside of their burrows, both times when they are most vulnerable. If prey can time these risky behaviors with periods of low fox activity or when foxes are absent from the area, their exposure to predation could be reduced. These dynamics could be incorporated by extending the current model to account for temporal variation in Arctic fox space use and activity, alongside prey risk-taking behavior.

### Perspectives

Predicting space use patterns of animals is a major challenge in ecology with many applications for the study of predator-prey dynamics (Suraci et al., 2022), conservation of connected areas (Hofmann et al., 2023), and the spread of invasive species (Forrest et al., 2025). Building mechanistic models of space use through step-selection functions rather than relying solely on correlative patterns should increase our ability to disentangle the mechanisms linking predators to their prey (Suraci et al., 2022). Such mechanistic models are also better suited to assess and predict how animals would react when exposed to novel or changing environments (Gomez et al., 2025). The development of predicting movement models of space use offers promising avenues to integrate other component of the predation sequence (Cherif et al., 2024). By building a predictive model of predator space-use, our study contributes to the development of more spatially informed approaches in predator–prey ecology.

## Acknowledgements

We thank Daniel Fortin, Dominique Gravel, Matthieu Weiss-Blais, Mathilde Poirier, Frédéric Letourneux, Alexandra Langwieder and David Bolduc for insightful discussions on the study. We are grateful to Marie-Christine Cadieux and Marie-Jeanne Rioux for their essential support in coordinating fieldwork campaigns and data management. We thank Mathieu Archambault, Laurianne Dumont, Marylou Beaudoin, Louis-Pierre Ouellet, Marie-Pier Poulin and the many people who collected data on Bylot Island for this project. We are grateful to the Sirmilik National Park of Canada and the community of Mittimatalik for their support. This work was financially supported by the Canada Foundation for Innovation, the Canada Research Chairs Program, the Kenneth M Molson Foundation, the Natural Sciences and Engineering Research Council of Canada (NSERC), the Canada First Sentinel North research program, the Network of Centres of Excellence of Canada ArcticNet, the Northern Scientific Training Program (Polar Knowledge Canada), and the Polar Continental Shelf Program (Natural Resources Canada). F. Dulude-de Broin received scholarships from NSERC, the Fonds de Recherche du Québec, and Sentinel North program.

## Author Contributions

Conceptualization: FDB, DB, JB, PL, Formal analysis: FDB (lead), CV (supporting), AGP (supporting), Data curation: FDB (lead), DB (supporting), AGP (supporting), JC (supporting), Visualization: FDB, Writing – original draft: FDB, Writing – review & editing: FDB, DB, JB, CV, AGP, AB, JC, AD, PL, Funding acquisition: DB, JB, AD, PL, Supervision: DB, JB, AD, PL.

## Data availability

Data and code required to reproduce the analyses will be made publicly available in an online repository upon acceptance.

## Supporting Information

### 1. SSF selection coefficients

**Figure 1a.**
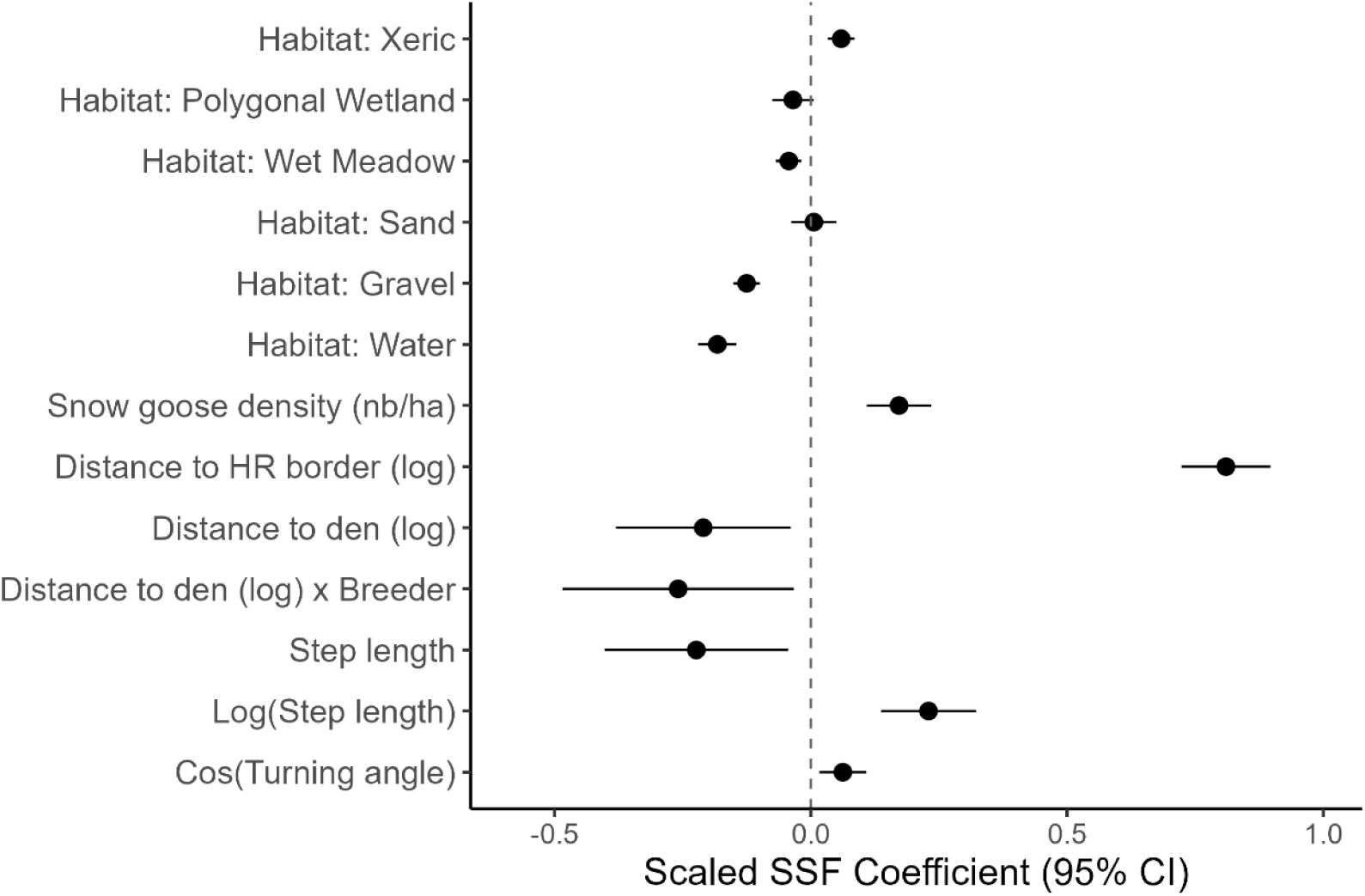
Selection coefficients of the step-selection function fitted to 4-minute relocation data collected from 31 individual-year foxes equipped with GPS collars on Bylot Island, Canada, between 2018–2023. All numeric variables were scaled before fitting the model and effect sizes are thus comparable. Dots represent SSF coefficient estimates and horizontal lines with their 95% confidence intervals.

### 2. Step-length and turning angle distributions from observed and simulated tracks

**Figure 7.**
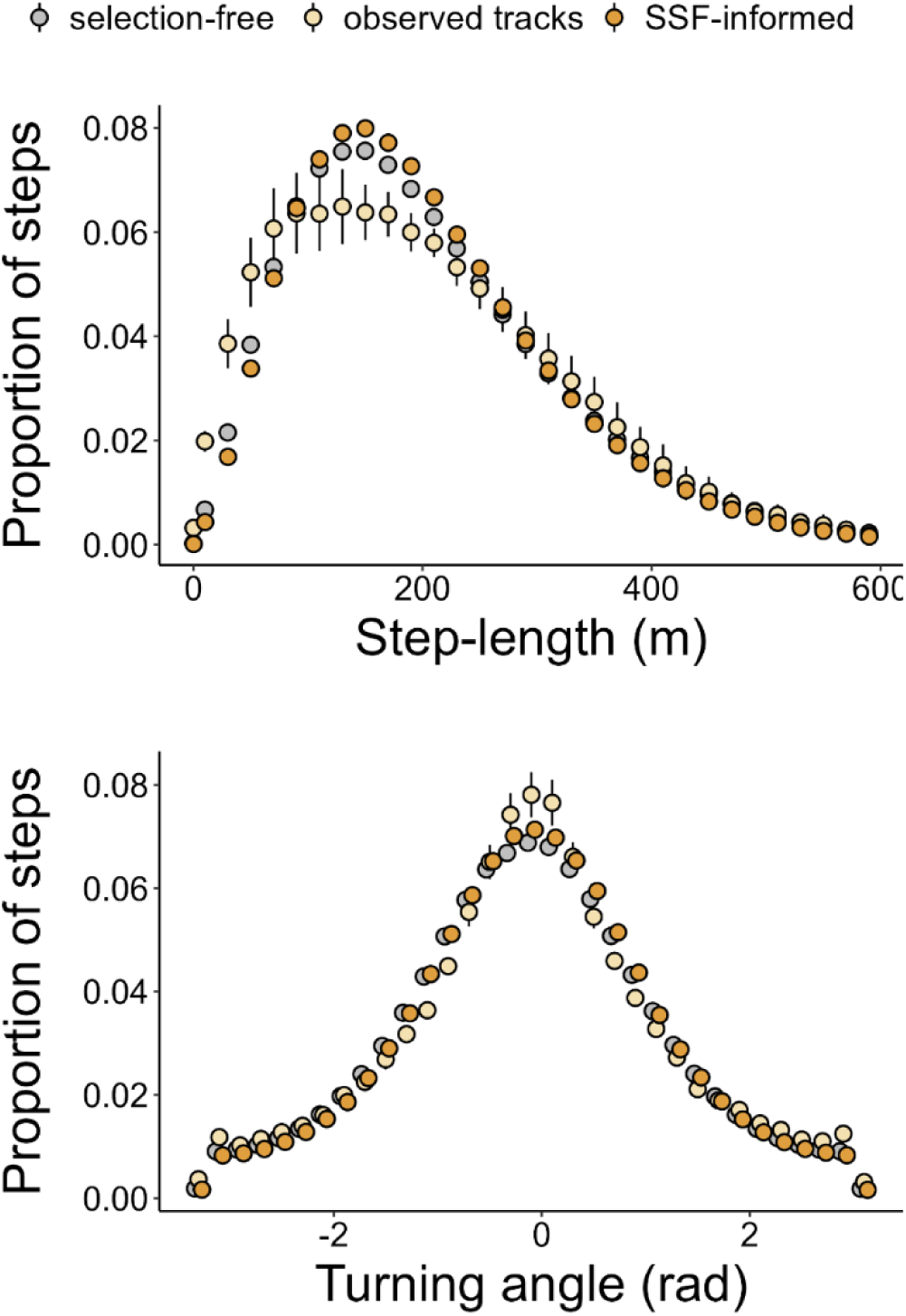
Comparison of step-length (top) and turning angles (bottom) distributions for observed tracks (yellow), simulated tracks generated from the SSF-informed movement model (orange) and simulated tracks generated from a selection-free movement model (gray). Dots represent the proportion of relocations within equally sized bins, averaged across all included individuals or simulations, with error bars indicating 95% confidence intervals.

### 3. Comparison of simulated and observed tracks on the restricted test sample

**Figure S1.**
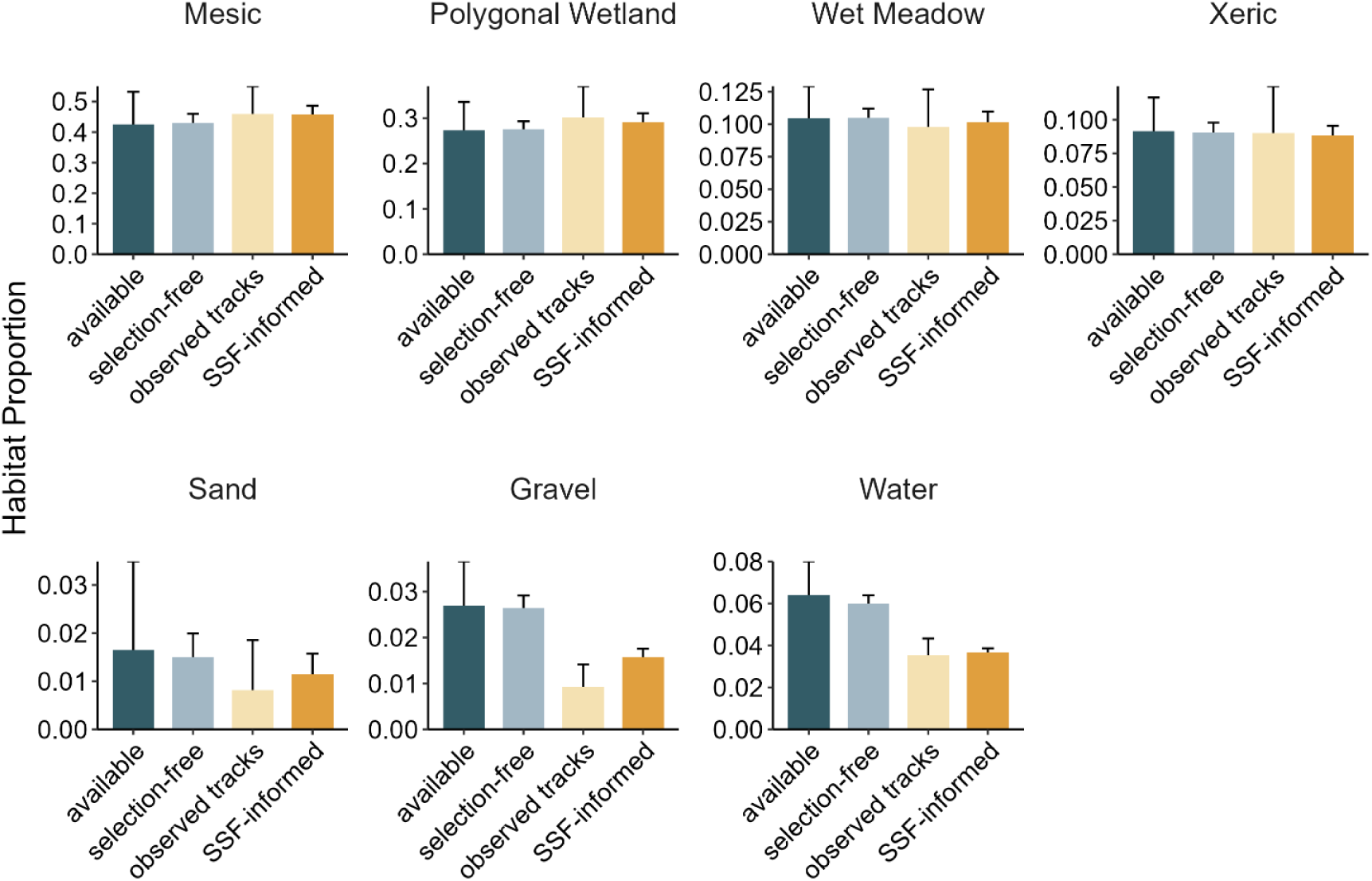
Comparison of average habitat class proportions from 8 test individuals (withheld when fitting the SSF model) across four categories: available (habitat available within fox home ranges), random (habitat visited by simulated tracks from a null movement model assuming no habitat selection), observed (habitat visited by GPS-collared foxes), and simulated (habitat visited by simulated tracks from the final movement model incorporating habitat selection). This comparison evaluates how closely habitat use by simulated fox tracks from the final model matches observed patterns on Bylot Island, Canada, between 2018 and 2023.

**Figure S2.**
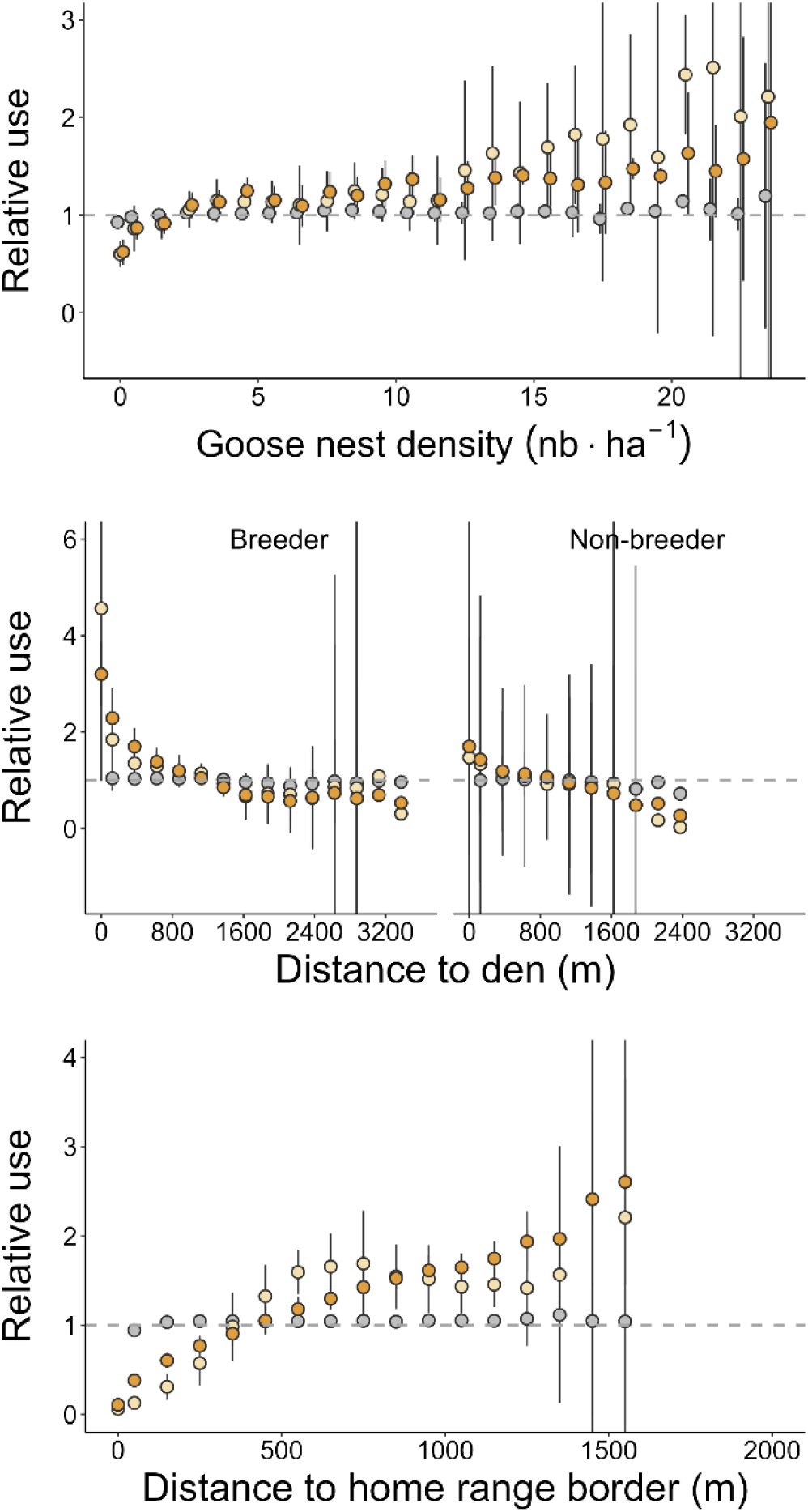
Comparison of the relative use of habitat patches with different goose densities, distance to den and distance to home range border for observed tracks (yellow), simulated tracks generated from the SSF-informed movement model (orange) and simulated tracks generated from a selection-free movement model (gray) for 8 test individuals withheld when fitting the SSF model. Dots represent the relative use (i.e. the ratio of used to available habitat patches) within equally sized bins, averaged across all included individuals or simulations, with error bars indicating 95% confidence intervals.

**Figure S3.**
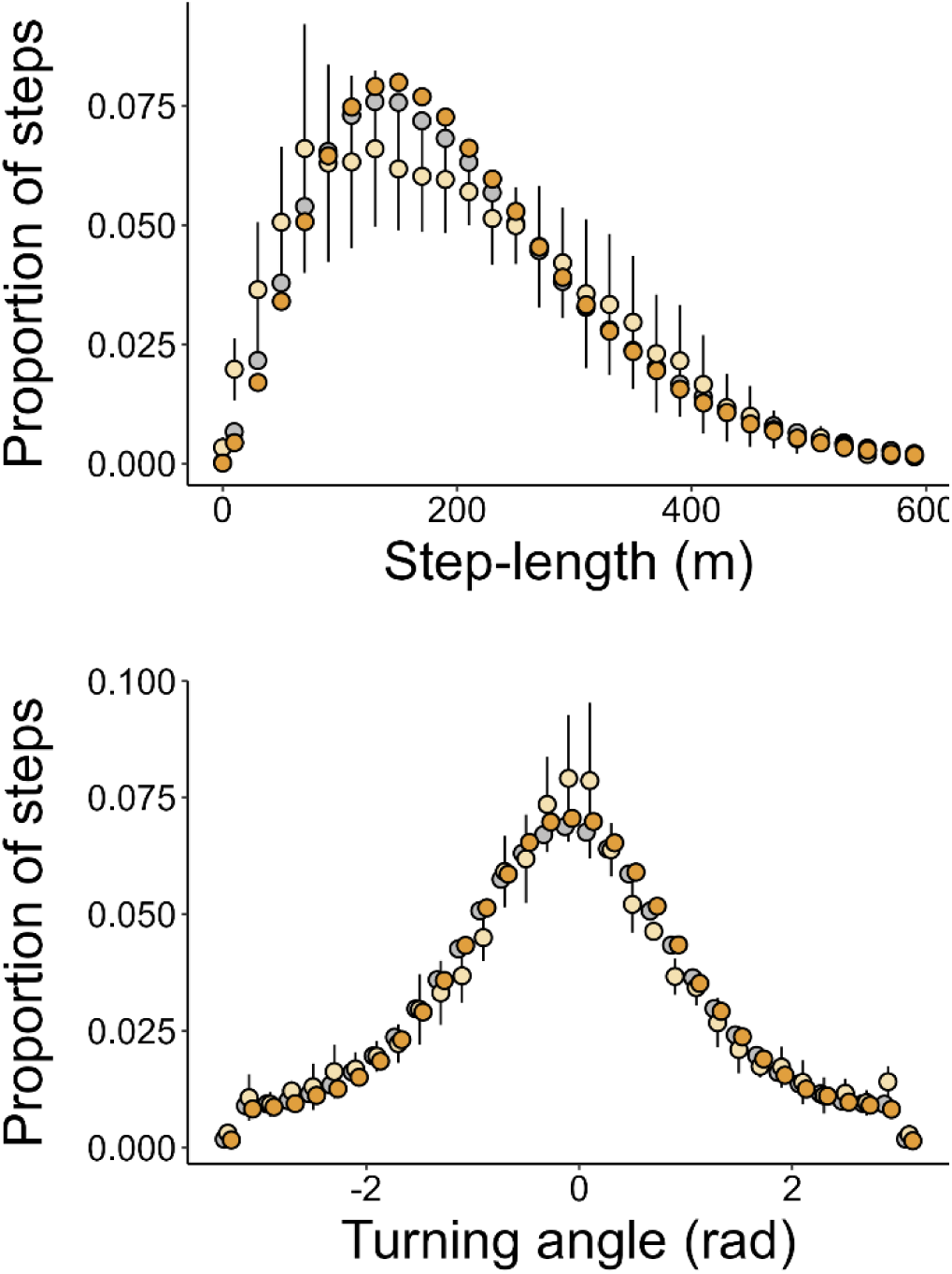
Comparison of step-length (top) and turning angles (bottom) distributions for observed tracks (yellow), simulated tracks generated from the SSF-informed movement model (orange) and simulated tracks generated from a selection-free movement model (gray) for 8 test individuals withheld when fitting the SSF model. Dots represent the proportion of relocations within equally sized bins, averaged across all included individuals or simulations, with error bars indicating 95% confidence intervals.

### 4. Comparison of relative habitat use with observed tracks across all individuals

**Table S1.** Results of linear mixed models comparing relative habitat use between observed and simulated fox tracks. The response variable was the ratio of used to available habitat patches, with data source (three-level factor: observed GPS-collared tracks, SSF-informed simulations, and selection-free simulations) as a fixed effect, and fox ID as a random effect. Confidence intervals excluding zero are highlighted in bold. Observed tracks served as the reference category, thus smaller coefficient estimates indicate closer similarity to observed fox movements, while confidence intervals excluding zero indicate significant differences from observed tracks.

|  | Estimate | 95%CI |  | Estimate | 95%CI |
| --- | --- | --- | --- | --- | --- |
| **Mesic habitat** |  |  | **Sand** |  |  |
| (Intercept) | **1.08** | **[1.06, 1.10]** | (Intercept) | **1.11** | **[0.57, 1.65]** |
| SSF-informed model | 0.00 | [-0.02, 0.01] | SSF-informed model | -0.13 | [-0.29, 0.04] |
| **Selection-free model** | **-0.07** | **[-0.09, -0.06]** | **Selection-free model** | **0.27** | **[0.11, 0.44]** |
| --- |  |  | --- |  |  |
| $\sigma$ fox id | 0.001 | | $\sigma$ fox id | 1.73 | |
| $\sigma$ residual | 0.002 | | $\sigma$ residual | 0.11 | |
| n = | 838 |  | n = | 442 |  |
| **Polygonal wetland** |  |  | **Gravel** |  |  |
| (Intercept) | **1.07** | **[1.04, 1.10]** | (Intercept) | **0.50** | **[0.44, 0.57]** |
| SSF-informed model | -0.04 | [-0.06, -0.01] | SSF-informed model | **0.15** | **[0.10, 0.20]** |
| **Selection-free model** | **-0.07** | **[-0.09, -0.04]** | **Selection-free model** | **0.47** | **[0.42, 0.52]** |
| --- |  |  | --- |  |  |
| $\sigma$ fox id | 0.002 | | $\sigma$ fox id | 0.017 | |
| $\sigma$ residual | 0.006 | | $\sigma$ residual | 0.024 | |
| n = | 838 |  | n = | 838 |  |
| **Wet meadow** |  |  | **Water** |  |  |
| (Intercept) | **0.95** | **[0.92, 0.98]** | (Intercept) | **0.58** | **[0.54, 0.62]** |
| SSF-informed model | **0.03** | **[0.01, 0.06]** | SSF-informed model | **0.04** | **[0.02, 0.07]** |
| **Selection-free model** | **0.05** | **[0.03, 0.08]** | **Selection-free model** | **0.39** | **[0.36, 0.42]** |
| --- |  |  | --- |  |  |
| $\sigma$ fox id | 0.003 | | $\sigma$ fox id | 0.008 | |
| $\sigma$ residual | 0.005 | | $\sigma$ residual | 0.008 | |
| n = | 838 |  | n = | 838 |  |
| **Snow goose nest density** |  |  | **Distance to den** |  |  |
| (Intercept) | **1.28** | **[1.23, 1.34]** | (Intercept) | **1.22** | **[1.15, 1.29]** |
| SSF-informed model | -0.15 | [-0.20, -0.11] | SSF-informed model | -0.09 | [-0.19, 0.01] |
| **Selection-free model** | **-0.29** | **[-0.33, -0.24]** | **Selection-free model** | **-0.23** | **[-0.33, -0.12]** |
| --- |  |  | --- |  |  |
| $\sigma$ fox id | 0.02 | | $\sigma$ fox id | 0.0004 | |
| $\sigma$ residual | 0.20 | | $\sigma$ residual | 0.62 | |
| n = | 2268 |  | n = | 1383 |  |
| **Distance to homerange border** |  |  |  |  |  |
| (Intercept) | **1.26** | **[1.20, 1.32]** |  |  |  |
| SSF-informed model | -0.03 | [-0.10, 0.05] |  |  |  |
| **Selection-free model** | **-0.25** | **[-0.32, -0.17]** |  |  |  |
| --- |  |  |  |  |  |
| $\sigma$ fox id | 0.004 | | | | |
| $\sigma$ residual | 0.45 | | | | |
| n = | 1737 |  |  |  |  |

### 1. Comparison of relative habitat use with observed tracks on test individuals only

**Table S2.** Results of linear mixed models comparing relative habitat use between observed and simulated fox tracks for 8 test individuals withheld when fitting the SSF model. The response variable was the ratio of used to available habitat patches, with data source (three-level factor: observed GPS-collared tracks, SSF-informed simulations, and selection-free simulations) as a fixed effect, and fox ID as a random effect. Confidence intervals excluding zero are highlighted in bold. Observed tracks served as the reference category, thus smaller coefficient estimates indicate closer similarity to observed fox movements, while confidence intervals excluding zero indicate significant differences from observed tracks.

|  | Estimate | 95%CI |  | Estimate | 95%CI |
| --- | --- | --- | --- | --- | --- |
| **Mesic habitat** |  |  | **Sand** |  |  |
| (Intercept) | **1.09** | **[1.04, 1.15]** | (Intercept) | **0.66** | **[0.15, 1.16]** |
| SSF-informed model | 0.00 | [-0.04, 0.05] | SSF-informed model | -0.05 | [-0.42, 0.32] |
| **Selection-free model** | **-0.08** | **[-0.12, -0.04]** | **Selection-free model** | **0.26** | **[-0.11, 0.63]** |
| --- |  |  | --- |  |  |
| $\sigma$ fox id | 0.003 | | $\sigma$ fox id | 0.20 | |
| $\sigma$ residual | 0.003 | | $\sigma$ residual | 0.16 | |
| n = | 157 |  | n = | 120 |  |
| **Polygonal wetland** |  |  | **Gravel** |  |  |
| (Intercept) | **1.08** | **[1.01, 1.14]** | (Intercept) | **0.41** | **[0.27, 0.54]** |
| SSF-informed model | -0.01 | [-0.07, 0.04] | **SSF-informed model** | **0.22** | **[0.12, 0.33]** |
| **Selection-free model** | **-0.07** | **[-0.13, -0.02]** | **Selection-free model** | **0.57** | **[0.47, 0.68]** |
| --- |  |  | --- |  |  |
| $\sigma$ fox id | 0.003 | | $\sigma$ fox id | 0.014 | |
| $\sigma$ residual | 0.005 | | $\sigma$ residual | 0.019 | |
| n = | 157 |  | n = | 157 |  |
| **Wet meadow** |  |  | **Water** |  |  |
| (Intercept) | **0.93** | **[0.88, 0.99]** | (Intercept) | **0.65** | **[0.57, 0.72]** |
| SSF-informed model | 0.03 | [-0.02, 0.08] | **SSF-informed model** | **-0.06** | **[-0.11, -0.01]** |
| **Selection-free model** | **0.07** | **[0.02, 0.12]** | **Selection-free model** | **0.30** | **[0.24, 0.35]** |
| --- |  |  | --- |  |  |
| $\sigma$ fox id | 0.002 | | $\sigma$ fox id | 0.005 | |
| $\sigma$ residual | 0.004 | | $\sigma$ residual | 0.005 | |
| n = | 157 |  | n = | 157 |  |
| **Snow goose nest density** |  |  | **Distance to den** |  |  |
| (Intercept) | **1.31** | **[1.22, 1.39]** | (Intercept) | **1.25** | **[1.05, 1.45]** |
| **SSF-informed model** | **-0.10** | **[-0.19, -0.01]** | SSF-informed model | -0.05 | [-0.30, 0.21] |
| **Selection-free model** | **-0.29** | **[-0.38, -0.20]** | Selection-free model | -0.25 | [-0.51, 0.01] |
| --- |  |  | --- |  |  |
| $\sigma$ fox id | 0.006 | | $\sigma$ fox id | 0.014 | |
| $\sigma$ residual | 0.15 | | $\sigma$ residual | 0.66 | |
| n = | 408 |  | n = | 231 |  |
| **Distance to homerange border** |  |  |  |  |  |
| (Intercept) | **1.17** | **[1.06, 1.28]** |  |  |  |
| SSF-informed model | 0.04 | [-0.11, 0.20] |  |  |  |
| **Selection-free model** | **-0.17** | **[-0.32, -0.01]** |  |  |  |
| --- |  |  |  |  |  |
| $\sigma$ fox id | 0.00 | | | | |
| $\sigma$ residual | 0.31 | | | | |
| n = | 306 |  |  |  |  |

## References

Aiello, C. M., Galloway, N. L., Prentice, P. R., Darby, N. W., Hughson, D., & Epps, C. W. (2023). Movement models and simulation reveal highway impacts and mitigation opportunities for a metapopulation-distributed species. Landscape Ecology, 38(4), 1085–1103. 10.1007/s10980-023-01600-6

Allan, B. M., Nimmo, D. G., Ierodiaconou, D., VanDerWal, J., Koh, L. P., & Ritchie, E. G. (2018). Futurecasting ecological research: the rise of technoecology. Ecosphere, 9(5). 10.1002/ecs2.2163

Avgar, T., Potts, J. R., Lewis, M. A., & Boyce, M. S. (2016). Integrated step selection analysis: Bridging the gap between resource selection and animal movement. Methods in Ecology and Evolution, 7(5), 619–630. 10.1111/2041-210X.12528

Beardsell, A., Berteaux, D., Dulude-De Broin, F., Gauthier, G., Clermont, J., Gravel, D., & Bêty, J. (2023). Predator-mediated interactions through changes in predator home range size can lead to local prey exclusion. Proceedings of the Royal Society B: Biological Sciences, 290, 20231154. 10.1098/rspb.2023.1154

Beardsell, A., Gravel, D., Berteaux, D., Gauthier, G., Clermont, J., Careau, V., Lecomte, N., Juhasz, C. C., Royer-Boutin, P., & Bêty, J. (2021). Derivation of Predator Functional Responses Using a Mechanistic Approach in a Natural System. Frontiers in Ecology and Evolution, 9, 1–12. 10.3389/fevo.2021.630944

Beardsell, A., Gravel, D., Clermont, J., Berteaux, D., Gauthier, G., & Bêty, J. (2022). A mechanistic model of functional response provides new insights into indirect interactions among arctic tundra prey. Ecology, 103(8), 1–16. 10.1002/ecy.3734

Beltran, R. S., Kilpatrick, A. M., Picardi, S., Abrahms, B., Barrile, G. M., Oestreich, W. K., Smith, J. A., Czapanskiy, M. F., Favilla, A. B., Reisinger, R. R., Kendall-Bar, J. M., Payne, A. R., Savoca, M. S., Palance, D. G., Andrzejaczek, S., Shen, D. M., Adachi, T., Costa, D. P., Storm, N. A., … Robinson, P. W. (2024). Maximizing biological insights from instruments attached to animals. Trends in Ecology and Evolution, 40(1), 37–46. 10.1016/j.tree.2024.09.009

Benhamou, S., & Courbin, N. (2023). Accounting for central place foraging constraints in habitat selection studies. Ecology, *November* 2022, 1–9. 10.1002/ecy.4134

Beyer, H. L., Gurarie, E., Börger, L., Panzacchi, M., Basille, M., Herfindal, I., Van Moorter, B., R. Lele, S., & Matthiopoulos, J. (2016). “You shall not pass!”: Quantifying barrier permeability and proximity avoidance by animals. Journal of Animal Ecology, 85(1), 43–53. 10.1111/1365-2656.12275

Brooks, Mollie, E., Kristensen, K., Benthem, Koen, J., V., Magnusson, A., Berg, Casper, W., Nielsen, A., Skaug, Hans, J., Mächler, M., & Bolker, Benjamin, M. (2017). glmmTMB Balances Speed and Flexibility Among Packages for Zero-inflated Generalized Linear Mixed Modeling. The R Journal, 9(2), 378. 10.32614/RJ-2017-066

Brown, J. L., & Orians, G. H. (1970). Spacing Patterns in Mobile Animals. Annual Review of Ecology and Systematics, 1(1), 239–262. 10.1146/annurev.es.01.110170.001323

Buskirk, S. W., & Millspaugh, J. J. (2006). Metrics for Studies of Resource Selection. Journal of Wildlife Management, 70(2), 358–366. http://doi.org/10.2193/0022-541x(2006)70[358:mfsors]2.0.co;2

Calabrese, J. M., Fleming, C. H., & Gurarie, E. (2016). Ctmm: an R Package for Analyzing Animal Relocation Data As a Continuous-Time Stochastic Process. Methods in Ecology and Evolution, 7(9), 1124–1132. 10.1111/2041-210X.12559

Cameron, C., Berteaux, D., & Dufresne, F. (2011). Spatial variation in food availability predicts extrapair paternity in the arctic fox. Behavioral Ecology, 22(6), 1364–1373. 10.1093/beheco/arr158

Chen, Z., Pasher, J., Duffe, J., & Behnamian, A. (2017). Mapping Arctic Coastal Ecosystems with High Resolution Optical Satellite Imagery Using a Hybrid Classification Approach. Canadian Journal of Remote Sensing, 43(6), 513–527. 10.1080/07038992.2017.1370367

Cherif, M., Brose, U., Hirt, M. R., Ryser, R., Silve, V., Albert, G., Arnott, R., Berti, E., Cirtwill, A., Dyer, A., Gauzens, B., Gupta, A., Ho, H. C., Portalier, S. M. J., Wain, D., & Wootton, K. (2024). The environment to the rescue: can physics help predict predator–prey interactions? Biological Reviews, 33. 10.1111/brv.13105

Clermont, J., Dulude-de Broin, F., Poulin, M. P., & Berteaux, D. (2025). Territoriality Modulates the Effect of Conspecific Encounters on the Foraging Behaviours of a Mammalian Predator. Ecology and Evolution, 15(3), 1–17. 10.1002/ece3.71058

Costa-Pereira, R., Moll, R. J., Jesmer, B. R., & Jetz, W. (2022). Animal tracking moves community ecology: Opportunities and challenges. Journal of Animal Ecology, 91(7), 1334–1344. 10.1111/1365-2656.13698

Dröge, E., Creel, S., Becker, M. S., & M’Soka, J. (2017). Risky times and risky places interact to affect prey behaviour. Nature Ecology and Evolution, 1(8), 1123– 1128. 10.1038/s41559-017-0220-9

Duchesne, É., Lamarre, J., Gauthier, G., Berteaux, D., Gravel, D., & Bêty, J. (2021). Variable strength of predator-mediated effects on species occurrence in an arctic terrestrial vertebrate community. Ecography, 1–13. 10.1111/ecog.05760

Dulude-de Broin, F., Clermont, J., Beardsell, A., Ouellet, L. P., Legagneux, P., Bêty, J., & Berteaux, D. (2023). Predator home range size mediates indirect interactions between prey species in an arctic vertebrate community. Journal of Animal Ecology, 92(12), 2373–2385. 10.1111/1365-2656.14017

Fagan, W. F., Lewis, M. A., Auger-Méthé, M., Avgar, T., Benhamou, S., Breed, G., Ladage, L., Schlägel, U. E., Tang, W. W., Papastamatiou, Y. P., Forester, J., & Mueller, T. (2013). Spatial memory and animal movement. Ecology Letters, 16(10), 1316–1329. 10.1111/ele.12165

Fauteux, D., Gauthier, G., & Berteaux, D. (2015). Seasonal demography of a cyclic lemming population in the Canadian Arctic. Journal of Animal Ecology, 84(5), 1412–1422. 10.1111/1365-2656.12385

Fieberg, J., Signer, J., Smith, B., & Avgar, T. (2020). A ‘how-to’ guide for interpreting parameters in Resource- and Step-selection analyses. *BioRxiv*, November, 2020.11.12.379834.

Fleming, C. H., Fagan, W. F., Mueller, T., Olson, K. A., Leimgruber, P., & Calabrese, J. M. (2015). Rigorous home range estimation with movement data: a new autocorrelated kernel density estimator. Ecology, 96(5), 1182–1188. 10.1890/14-2010.1

Forrest, S. W., Pagendam, D., Bode, M., Drovandi, C., Potts, J. R., Perry, J., Vanderduys, E., & Hoskins, A. J. (2025). Predicting fine-scale distributions and emergent spatiotemporal patterns from temporally dynamic step selection simulations. Ecography, 2025(2), 1–16. 10.1111/ecog.07421

Fortin, D., Beyer, H. L., Boyce, M. S., Smith, D. W., Duchesne, T., & Mao, J. S. (2005). Wolves influence Elk movement: behaviour shapes a trophic cascade in Yellowstone National Park. Ecology, 86(5), 1320–1330.

Fortin, D., Buono, P. L., Schmitz, O. J., Courbin, N., Losier, C., St-Laurent, M. H., Drapeau, P., Heppell, S., Dussault, C., Brodeur, V., & Mainguy, J. (2015). A spatial theory for characterizing predator – Multiprey interactions in heterogeneous landscapes. Proceedings of the Royal Society B: Biological Sciences, 282(1812). 10.1098/rspb.2015.0973

Fretwell, S. D., & Lucas, H. L. (1969). On territorial behaviour and other factors influencing habitat distribution in birds. I. Theoretical Development. Acta Biotheoretica, 19, 16–36. 10.1039/9781847558213-00059

Gauthier, G., Berteaux, D., Bêty, J., Tarroux, A., Therrien, J.-F., McKinnon, L., Legagneux, P., & Cadieux, M.-C. (2011). The tundra food web of Bylot Island in a changing climate and the role of exchanges between ecosystems. Écoscience, 18(3), 223–235. 10.2980/18-3-3453

Gehr, B., Hofer, E. J., Ryser, A., Vimercati, E., Vogt, K., & Keller, L. F. (2018). Evidence for nonconsumptive effects from a large predator in an ungulate prey? Behavioral Ecology, 29(3), 724–735. 10.1093/beheco/ary031

Giroux, M. A., Berteaux, D., Lecomte, N., Gauthier, G., Szor, G., & Bêty, J. (2012). Benefiting from a migratory prey: Spatio-temporal patterns in allochthonous subsidization of an arctic predator. Journal of Animal Ecology, 81(3), 533–542. 10.1111/j.1365-2656.2011.01944.x

Gomez, S., English, H. M., Bejarano, V., Paul, A., Bracken, A. M., Bray, E., Evans, L. C., Gan, J. L., Grecian, W. J., Gutmann, C., Seth, R., Hejcmanová, P., Lelotte, L., Michael, B., Potts, J. R., Russell, C. J. G., Rutz, C., Singh, N. J., Whyte, K. F., … Gomez, S. (2025). Understanding and predicting animal movements and distributions in the Anthropocene. March, 1–19. 10.1111/1365-2656.70040

Grenier-Potvin, A., Clermont, J., Gauthier, G., & Berteaux, D. (2021). Prey and habitat distribution are not enough to explain predator habitat selection: addressing intraspecific interactions, behavioural state and time. Movement Ecology, 9(1), 1–13. 10.1186/s40462-021-00250-0

Gruyer, N., Gauthier, G., & Berteaux, D. (2008). Cyclic dynamics of sympatric lemming populations on Bylot Island, Nunavut, Canada. Canadian Journal of Zoology, 86(8), 910–917. 10.1139/Z08-059

Heithaus, M. R., Wirsing, A. J., Burkholder, D., Thomson, J., & Dill, L. M. (2009). Towards a predictive framework for predator risk effects: The interaction of landscape features and prey escape tactics. Journal of Animal Ecology, 78(3), 556–562. 10.1111/j.1365-2656.2008.01512.x

Hertel, A. G., Hertel, A. G., Niemelä, P. T., Dingemanse, N. J., Mueller, T., & Mueller, T. (2020). A guide for studying among-individual behavioral variation from movement data in the wild. Movement Ecology, 8(1), 1–18. 10.1186/s40462-020-00216-8

Hofmann, D. D., Cozzi, G., McNutt, J. W., Ozgul, A., & Behr, D. M. (2023). A three-step approach for assessing landscape connectivity via simulated dispersal: African wild dog case study. Landscape Ecology, 38(4), 981–998. 10.1007/s10980-023-01602-4

Jeltsch, F., Bonte, D., Pe’er, G., Reineking, B., Leimgruber, P., Balkenhol, N., Schröder, B., Buchmann, C. M., Mueller, T., Blaum, N., Zurell, D., Böhning-Gaese, K., Wiegand, T., Eccard, J. A., Hofer, H., Reeg, J., Eggers, U., & Bauer, S. (2013). Integrating movement ecology with biodiversity research - exploring new avenues to address spatiotemporal biodiversity dynamics. Movement Ecology, 1(1), 1–13. 10.1186/2051-3933-1-6

Kays, R., Crofoot, M. C., Jetz, W., & Wikelski, M. (2015). Terrestrial animal tracking as an eye on life and planet. Science, 348(6240), aaa2478. 10.1126/science.aaa2478

Labadie, G., Hardy, C., Boulanger, Y., Vanlandeghem, V., Hebblewhite, M., & Fortin, D. (2023). Global change risks a threatened species due to alteration of predator–prey dynamics. Ecosphere, 14(3), 1–19. 10.1002/ecs2.4485

LaBarge, L. R., Krofel, M., Allen, M. L., Hill, R. A., Welch, A. J., & Allan, A. T. L. (2024). Keystone individuals – linking predator traits to community ecology. Trends in Ecology and Evolution, 39(11), 983–994. 10.1016/j.tree.2024.07.001

Léandri-Breton, D. J., & Bêty, J. (2020). Vulnerability to predation may affect species distribution: plovers with broader arctic breeding range nest in safer habitat. Scientific Reports, 10(1), 1–8. 10.1038/s41598-020-61956-6

Lecomte, N., Careau, V., Gauthier, G., & Giroux, J.-F. (2008). Predator behaviour and predation risk in the heterogeneous Arctic environment. Journal of Animal Ecology, 77(3), 439–447. 10.1111/j.1365-2656.2008.01354.x

Lichtenstein, J. L. L., Daniel, K. A., Wong, J. B., Wright, C. M., Doering, G. N., Costa-Pereira, R., & Pruitt, J. N. (2019). Habitat structure changes the relationships between predator behavior, prey behavior, and prey survival rates. Oecologia, 190(2), 297–308. 10.1007/s00442-019-04344-w

López-Sepulcre, A., & Kokko, H. (2005). Territorial Defense, Territory Size, and Population Regulation. The American Naturalist, 166(3), 317–328.

Loveridge, A. J., Valeix, M., Davidson, Z., Murindagomo, F., Fritz, H., & MacDonald, D. W. (2009). Changes in home range size of African lions in relation to pride size and prey biomass in a semi-arid savanna. Ecography, 32(6), 953–962. 10.1111/j.1600-0587.2009.05745.x

Manly. (2002). Resource Selection by Animals Statistical Design and Analysis for Field Studies Second Edition. Kluwer, Norwell, Mass, USA, 65(3), 25–28.

Mckinnon, L., Berteaux, D., Gauthier, G., & Bêty, J. (2013). Predator-mediated interactions between preferred, alternative and incidental prey in the arctic tundra. Oikos, 122(7), 1042–1048. 10.1111/j.1600-0706.2012.20708.x

Michelot, T., Langrock, R., & Patterson, T. (2018). moveHMM An R package for the analysis of animal movement data. ArXiv Preprint, 1–24.

Moorcroft, P. R., Barnett, A., Moorcroft, P. R., & Barnett, A. (2017). Mechanistic Home Range Models and Resource Selection Analysis : A Reconciliation and Unification Published by : Wiley on behalf of the Ecological Society of America Stable URL : http://www.jstor.org/stable/27651649 *REFERENCES Linked references are availab*. 89(4), 1112–1119.

Muff, S., Signer, J., & Fieberg, J. (2020). Accounting for individual-specific variation in habitat-selection studies: Efficient estimation of mixed-effects models using Bayesian or frequentist computation. Journal of Animal Ecology, 89(1), 80–92. 10.1111/1365-2656.13087

Nathan, R., Getz, W. M., Revilla, E., Holyoak, M., Kadmon, R., Saltz, D., & Smouse, P. E. (2008). A movement ecology paradigm for unifying organismal movement research. Proceedings of the National Academy of Sciences, 105(49), 19052– 19059. 10.1073/pnas.0800375105

Nathan, R., Monk, C. T., Arlinghaus, R., Adam, T., Alós, J., Assaf, M., Baktoft, H., Beardsworth, C. E., Bertram, M. G., Bijleveld, A. I., Brodin, T., Brooks, J. L., Campos-Candela, A., Cooke, S. J., Gjelland, K., Gupte, P. R., Harel, R., Hellström, G., Jeltsch, F., … Jarić, I. (2022). Big-data approaches lead to an increased understanding of the ecology of animal movement. Science, 375(6582). 10.1126/science.abg1780

Neumann, W., Martinuzzi, S., Estes, A. B., Pidgeon, A. M., Dettki, H., Ericsson, G., & Radeloff, V. C. (2015). Opportunities for the application of advanced remotely-sensed data in ecological studies of terrestrial animal movement. Movement Ecology, 3(1), 1–13. 10.1186/s40462-015-0036-7

Oliver, M., Jose, J., & Lambin, X. (2009). Do rabbits eat voles *?* Apparent competition, habitat heterogeneity and large-scale coexistence under mink predation. 1201–1209. 10.1111/j.1461-0248.2009.01375.x

Palmer, M. S., Gaynor, K. M., Becker, J. A., Abraham, J. O., Mumma, M. A., & Pringle, R. M. (2022). Dynamic landscapes of fear: understanding spatiotemporal risk. Trends in Ecology & Evolution, 37(10), 911–925. 10.1016/j.tree.2022.06.007

Patterson, T. A., Thomas, L., Wilcox, C., Ovaskainen, O., & Matthiopoulos, J. (2008). State-space models of individual animal movement. Trends in Ecology and Evolution, 23(2), 87–94. 10.1016/j.tree.2007.10.009

Potts, J. R., Auger-Méthé, M., Mokross, K., & Lewis, M. A. (2014). A generalized residual technique for analysing complex movement models using earth mover’s distance. Methods in Ecology and Evolution, 5(10), 1012–1022. 10.1111/2041-210X.12253

Potts, J. R., Bastille-Rousseau, G., Murray, D. L., Schaefer, J. A., & Lewis, M. A. (2014). Predicting local and non-local effects of resources on animal space use using a mechanistic step selection model. Methods in Ecology and Evolution, 5(3), 253–262. 10.1111/2041-210X.12150

Potts, J. R., & Börger, L. (2023). How to scale up from animal movement decisions to spatiotemporal patterns: An approach via step selection. Journal of Animal Ecology, 92(1), 16–29. 10.1111/1365-2656.13832

Potts, J. R., & Lewis, M. A. (2014). How do animal territories form and change? lessons from 20 years of mechanistic modelling. Proceedings of the Royal Society B: Biological Sciences, 281(1784). 10.1098/rspb.2014.0231

Reed, A., Hughes, R. J., & Boyd, H. (2002). Patterns of distribution and abundance of Greater Snow Geese on Bylot Island, Nunavut, Canada 1983-1998. Wildfowl, 53, 53–65.

Schmitz, O. J., Miller, J. R. B., Trainor, A. M., & Abrahms, B. (2017). Toward a community ecology of landscapes: predicting multiple predator--prey interactions across geographic space. Ecology, 98(9), 2281–2292. 10.1002/ecy.1916

Schuhmacher, D., Bähre, B., Bonneel, N., Gottschlich, Carsten Hartmann, V., Heinemann, F., Schrieber, B., & Schmitzer, J. (2024). transport: Computation of Optimal Transport Plans and Wasserstein Distances. R Package Version 0.15-4.

Sells, S. N., & Mitchell, M. S. (2020). The economics of territory selection. Ecological Modelling, 438(October), 109329. 10.1016/j.ecolmodel.2020.109329

Sells, S. N., Mitchell, M. S., Ausband, D. E., Luis, A. D., Emlen, D. J., Podruzny, K. M., & Gude, J. A. (2022). Economical defence of resources structures territorial space use in a cooperative carnivore. Proceedings of the Royal Society B: Biological Sciences, 289, 20212512. 10.1098/rspb.2021.2512

Signer, J., Fieberg, J., Reineking, B., Schlägel, U., Smith, B., Balkenhol, N., & Avgar, T. (2024). Simulating animal space use from fitted integrated Step-Selection Functions (iSSF). Methods in Ecology and Evolution, 15(1), 43–50. 10.1111/2041-210X.14263

Sih, A. (2005). Predator-prey space use as an emergent outcome of a behavioral response race. In Ecology of predator-prey interactions (Vol. 256, p. 78). Oxford university press New York.

Smith, J. A., Donadio, E., Pauli, J. N., Sheriff, M. J., Bidder, O. R., & Middleton, A. D. (2019). Habitat complexity mediates the predator–prey space race. Ecology, 100(7), 1–9. 10.1002/ecy.2724

Smith, J. A., Donadio, E., Pauli, J. N., Sheriff, M. J., & Middleton, A. D. (2019). Integrating temporal refugia into landscapes of fear: prey exploit predator downtimes to forage in risky places. Oecologia, 189(4), 883–890. 10.1007/s00442-019-04381-5

Smouse, P. E., Focardi, S., Moorcroft, P. R., Kie, J. G., Forester, J. D., & Morales, J. M. (2010). Stochastic modelling of animal movement. Philosophical Transactions of the Royal Society B: Biological Sciences, 365(1550), 2201– 2211. 10.1098/rstb.2010.0078

Stephens, D. W., & Krebs, J. (1986). Foraging theory. Princeton university press.

Suraci, J. P., Smith, J. A., Chamaillé-Jammes, S., Gaynor, K. M., Jones, M., Luttbeg, B., Ritchie, E. G., Sheriff, M. J., & Sih, A. (2022). Beyond spatial overlap: harnessing new technologies to resolve the complexities of predator–prey interactions. Oikos, 2022(8), 1–15. 10.1111/oik.09004

Szor, G., Berteaux, D., & Gauthier, G. (2008). Finding the right home: Distribution of food resources and terrain characteristics influence selection of denning sites and reproductive dens in arctic foxes. Polar Biology, 31(3), 351–362. 10.1007/s00300-007-0364-1

Toscano, B. J., Gownaris, N. J., Heerhartz, S. M., & Monaco, C. J. (2016). Personality, foraging behavior and specialization: integrating behavioral and food web ecology at the individual level. Oecologia, 182(1), 55–69. 10.1007/s00442-016-3648-8

Toscano, B. J., & Griffen, B. D. (2014). Trait-mediated functional responses: Predator behavioural type mediates prey consumption. Journal of Animal Ecology, 83(6), 1469–1477. 10.1111/1365-2656.12236

Vanlandeghem, V., Drapeau, P., Prima, M. C., St-Laurent, M. H., & Fortin, D. (2021). Management-mediated predation rate in the caribou–moose–wolf system: spatial configuration of logging activities matters. Ecosphere, 12(6). 10.1002/ecs2.3550

Williams, H. J., Taylor, L. A., Benhamou, S., Bijleveld, A. I., Clay, T. A., de Grissac, S., Demšar, U., English, H. M., Franconi, N., Gómez-Laich, A., Griffiths, R. C., Kay, W. P., Morales, J. M., Potts, J. R., Rogerson, K. F., Rutz, C., Spelt, A., Trevail, A. M., Wilson, R. P., & Börger, L. (2020). Optimizing the use of biologgers for movement ecology research. Journal of Animal Ecology, 89(1), 186–206. 10.1111/1365-2656.13094

Wilmers, C. C., Nickel, B., Bryce, C. M., Smith, J. A., Wheat, R. E., Yovovich, V., & Hebblewhite, M. (2015). The golden age of bio-logging: How animal-borne sensors are advancing the frontiers of ecology. Ecology, 96(7), 1741–1753. 10.1890/14-1401.1

Winter, V. A., Smith, B. J., Berger, D. J., Hart, R. B., Huang, J., Manlove, K., Buderman, F. E., & Avgar, T. (2024). Forecasting animal distribution through individual habitat selection: insights for population inference and transferable predictions. Ecography, 1–15. 10.1111/ecog.07225

Wootton, K. L., Curtsdotter, A., Roslin, T., Bommarco, R., & Jonsson, T. (2023). Towards a modular theory of trophic interactions. Functional Ecology, 37(1), 26–43. 10.1111/1365-2435.13954

Zurell, D., König, C., Malchow, A. K., Kapitza, S., Bocedi, G., Travis, J., & Fandos, G. (2022). Spatially explicit models for decision-making in animal conservation and restoration. Ecography, 2022(4), 1–16. 10.1111/ecog.05787

